# Age-driven miR-29 trades youthful plasticity for epigenetic fidelity

**DOI:** 10.64898/2026.10.18.759616

**Authors:** Ilaria Craparotta, Maria Chiara Barbera, Giada Andrea Cassanmagnago, Anna Chiara Salvatore, Alisia Sallemi, Luca Guarrera, Arianna Vallerga, Marco Bolis

**Affiliations:** Laboratory of Integrative Biomedicine, Department of Oncology, Istituto di Ricerche Farmacologiche ‘Mario Negri’ IRCCS, Via Mario Negri 2, 20156 Milano, Italy; Department of Biotechnology and Biosciences, University of Milano-Bicocca, Milano, Italy; Bioinformatics Core Unit, Institute of Oncology Research (IOR), Bellinzona, TI 6500, Switzerland; Università Della Svizzera Italiana (USI), Faculty of Biomedical Sciences, Bellinzona, TI 6500, Switzerland; Swiss Institute of Bioinformatics, Bioinformatics Core Unit, Bellinzona, TI 6500, Switzerland; Department of Informatics, Systems and Communication, University of Milano-Bicocca, Milano, Italy

## Abstract

Throughout life, somatic cells must preserve their identity while retaining the capacity to adapt. How this balance shifts with ageing and rejuvenation remains unclear. Here we identify miR-29 as a central point of convergence between the transcriptional programmes of physiological ageing and two routes to rejuvenation: induced reprogramming and the natural reset of early development. miR-29 rises with ageing, falls in both rejuvenation settings and shows the most conserved age-associated increase across vertebrates. miR-29 inhibition opposes transcriptional ageing, whereas mimics promote an older transcriptional state. Methylome analysis reveals a broader function: inhibition releases the miR-29 brake on remodelling, and mimic treatment reinforces it. This release increases methylation at repeat-associated CpGs and lowers epigenetic-clock estimates through unequal contributions in opposite directions. Inhibition also amplifies stimulus-induced methylation changes but reduces fidelity to lineage-specific methylation patterns. We extend this relationship to cancer, where low miR-29 expression marks tumours with younger epigenetic-age estimates yet greater methylation divergence from their tissue of origin. These findings identify miR-29 as a control point linking ageing, rejuvenation and cancer, and support a maladaptive trade-off model in which epigenetic fidelity is preserved at the expense of the plasticity needed for methylome maintenance and adaptation.

---

Ageing changes the molecular state of a cell without necessarily changing its identity. Rejuvenation therefore presents a particular challenge: how can youthful molecular features be restored while preserving the programmes that sustain specialised cell function? At the epigenetic level, this challenge involves reconciling plasticity, the capacity for methylome remodelling, with fidelity to the methylation patterns characteristic of an established somatic state. Preserving a functional methylome requires active maintenance, not simply resistance to change ^1^. Epigenetic alterations are prominent features of ageing, and DNA-methylation clocks capture reproducible components of this process ^2,3^. These clocks combine methylation measurements across many sites to estimate DNA-methylation age (DNAmAge). A lower estimate can reflect changes that consistently point towards a younger age, or a balance in which age-lowering contributions outweigh age-increasing ones.

Reprogramming demonstrates that molecular age is not fixed. Partial reprogramming can reverse age-associated features without completing the transition to pluripotency, while early development provides a physiological setting for epigenetic-age resetting ^4,5,6^. How access to younger molecular states is regulated in differentiated cells remains unclear. Could mechanisms that preserve epigenetic stability also restrict the plasticity needed for rejuvenation?

We approached this question without selecting a candidate gene or pathway. We first intersected transcriptional programmes that rise during normal ageing with programmes repressed during two rejuvenation-associated transitions. This intersection identified a single shared transcript, *LINC-PINT*, which encompasses the miR-29a/b1 cluster. We therefore examined mature miR-29 family members across tissues and species. To test whether miR-29 actively shapes, rather than merely marks, age-associated cellular states, we perturbed the family in adult primary and independently sourced immortalized human fibroblasts, without introducing reprogramming factors, and measured transcriptional and DNA-methylation responses over time. Inhibition lowered transcriptional and epigenetic-age estimates but increased methylation divergence from a fibroblast reference, whereas mimic treatment promoted an older transcriptional state and constrained methylome remodelling. In cancer, where miR-29 is frequently downregulated ^7^ and cellular plasticity is often deregulated ^8^, we asked whether lower miR-29 was associated with greater divergence from normal-tissue methylation profiles. Together, these findings show that miR-29 inhibition reactivates youthful transcriptional programmes while releasing constraints on methylome remodelling. This renewed plasticity, however, can come at the cost of lineage-specific epigenetic fidelity.

## Ageing and rejuvenation converge on miR-29

We began with a human somatic-cell reprogramming time course in which cells progressing towards pluripotency (SSEA4-positive) could be separated from cells retaining a fibroblast-like state (CD13-positive). In the SSEA4-positive branch, DNAmAge fell sharply between approximately days 11 and 15; the CD13-positive branch retained an adult-like clock profile (Fig. 1a). Comparing transcription between these branches over days 11–15 identified 531 strongly repressed genes during reprogramming (programme R; Fig. 1b). This focused the analysis on the interval of DNAmAge decline rather than the entire reprogramming trajectory.

**Fig. 1.**
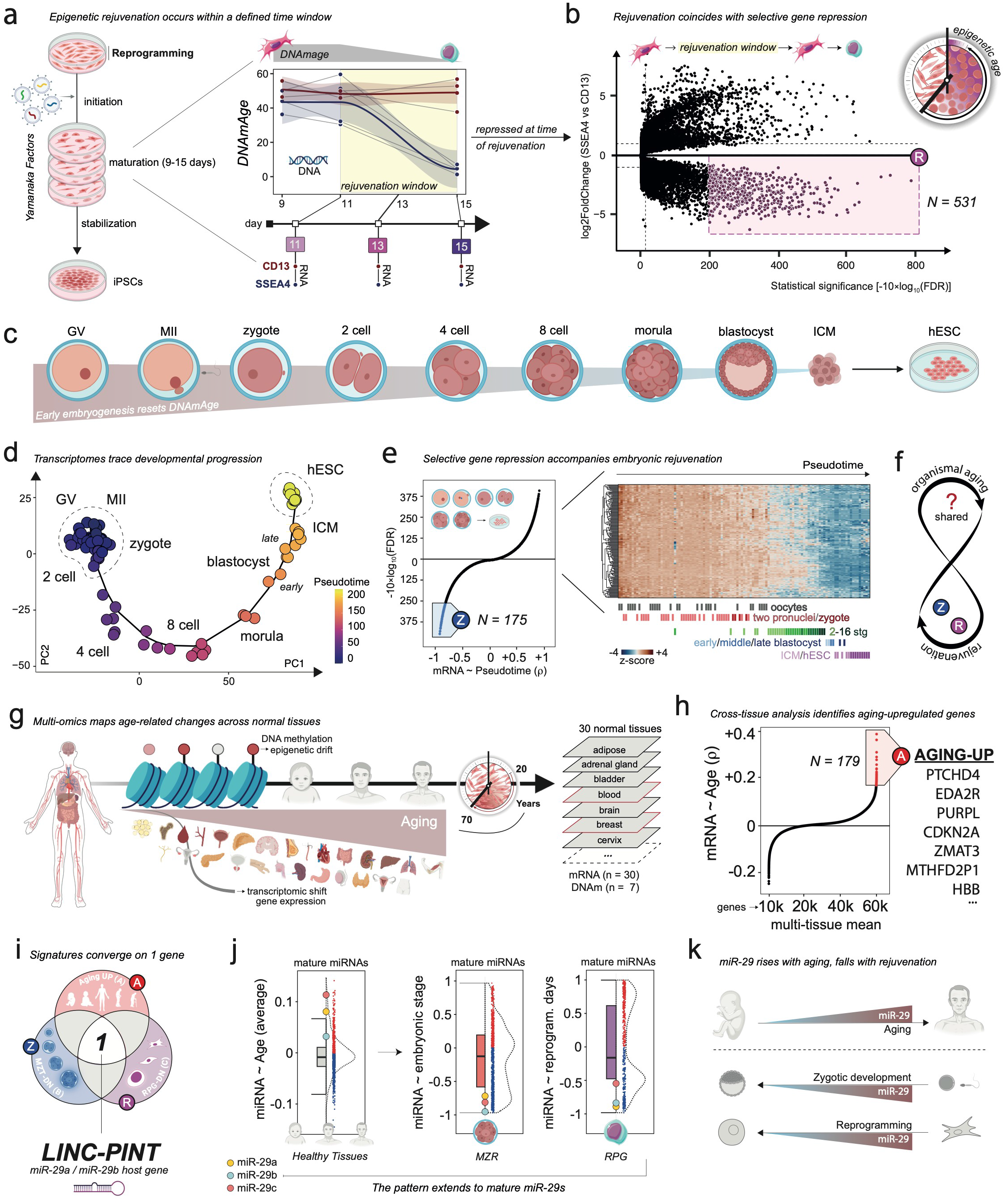
**Independent ageing and rejuvenation programmes converge on a miR-29-associated locus**. **a**, Reprogramming and Horvath DNA methylation age (DNAmAge) in SSEA4-positive and CD13-positive cells: 22 methylation profiles from four adult donors. Grey lines connect donor branches; LOESS fits have 90% confidence bands. DNAmAge is shown at days 9, 11 and 15; RNA-seq focuses on days 11, 13 and 15, spanning clock decline. **b**, Branch expression differences over days 11–15, adjusted for donor and time (36 RNA profiles; six donors, including two neonatal donors). The negative three-standard-deviation tail of DESeq2 Wald statistics defines reprogramming-down signature R (531 genes). c,d, Early-development schematic, principal-component projection and pseudotime across five RNA-seq cohorts spanning oocytes to human embryonic stem cells. *Age-driven miR-29 trades youthful plasticity for epigenetic fidelity* **e**, Developmental-down signature Z (175 genes): pseudotime correlations and row-standardised expression. Selection uses the negative three-standard-deviation tail of Fisher-transformed Spearman coefficients, separately from the signed significance axis. **f**, Framework for identifying transcripts increasing with ageing and decreasing during rejuvenation. **g**, Normal-tissue RNA and methylation resources: 30 RNA-screen tissue categories and seven paired-analysis tissues. **h**, Cross-tissue chronological-age ranking; its positive three-standard-deviation tail defines signature A (179 genes). **i**, The A/R/Z intersection identifies *LINC-PINT*, a miR-29a/b1 host gene, as the sole shared transcript. **j**, Mature miR-29a, miR-29b and miR-29c within miRNA correlations with age, embryonic stage and reprogramming time. Boxes show medians and interquartile ranges of miRNA coefficients; whiskers extend to 1.5 interquartile ranges. **k**, Reciprocal ageing and rejuvenation pattern. Source datasets and aggregation are described in Methods.

Early embryogenesis provides a physiological context of epigenetic rejuvenation ^6^. We therefore reconstructed the developmental transcriptional trajectory from oocytes through early embryonic and pluripotent states, ordering RNA profiles by pseudotime (Fig. 1c,d). We identified 175 genes whose expression declined strongly along this trajectory (programme Z; Fig. 1e). We then asked which transcripts repressed in both settings also increased during normal ageing (Fig. 1f).

The ageing comparison spanned 30 normal human tissue categories, with a smaller matched RNA– methylation resource for comparisons with DNAmAge (Fig. 1g and Supplementary Fig. 1a–d) ^9^. The cross-tissue ranking identified 179 genes with strong positive associations with age (programme A; Fig. 1h). Each programme was defined within its own dataset. *LINC-PINT* was the only transcript shared by the three programmes, increasing with ageing (A) and decreasing in both rejuvenation-associated programmes (R and Z) (Fig. 1i). A complementary integration of continuous directional ranks, without tail cutoffs, also placed *LINC-PINT* first among 34,686 jointly assessed genes (Supplementary Methods).

Across the seven tissues profiled for both gene expression and DNA methylation, *LINC-PINT* ranked higher when expression was related to DNAmAge rather than chronological age (Supplementary Fig. 1e). The positive chronological-age association also persisted after donor aggregation (Supplementary Fig. 1f). In strictly matched RNA–methylation samples, its expression was positively associated with both DNAmAge and age acceleration (Supplementary Fig. 1g).

*LINC-PINT*, the sole transcript shared by A, R and Z (Supplementary Fig. 1h), is annotated as a host gene for the miR-29a/b1 cluster. A long primary transcript spanning this cluster shares the *LINC-PINT* transcription start site, whereas shorter miRNA transcripts initiate independently ^10^. We therefore asked whether the signal extended to mature miRNAs. miR-29 family members occupied the age-positive side of the healthy-tissue distributions and showed the opposite behaviour during reprogramming and early development (Fig. 1j, Supplementary Fig. 1i). The result extended to miR-29c, encoded at the second human miR-29 locus. Thus, a transcriptome-first intersection led to a family-level signal linking ageing and rejuvenation-associated transitions (Fig. 1k).

## A conserved age-associated signal across vertebrates

We next asked whether the age-associated increase in miR-29 observed in humans reflects a broadly conserved feature of molecular ageing across species with markedly different lifespans. To make these comparisons, we first reconstructed the evolutionary relationships among miR-29 family members. Our reconstruction traced miR-29 back to early bilaterian evolution (Fig. 2a). We then focused on vertebrates, where counterparts of the two human genomic clusters remain recognisable despite lineage-specific diversification (Fig. 2a,b). The human family contains four genomic miRNA copies that produce three mature forms, miR-29a, miR-29b and miR-29c, with a shared seed (Fig. 2c). Sequence and locus reconstruction resolved the relationships among these family members across vertebrates (Fig. 2d,e, Supplementary Fig. 2a).

**Fig. 2.**
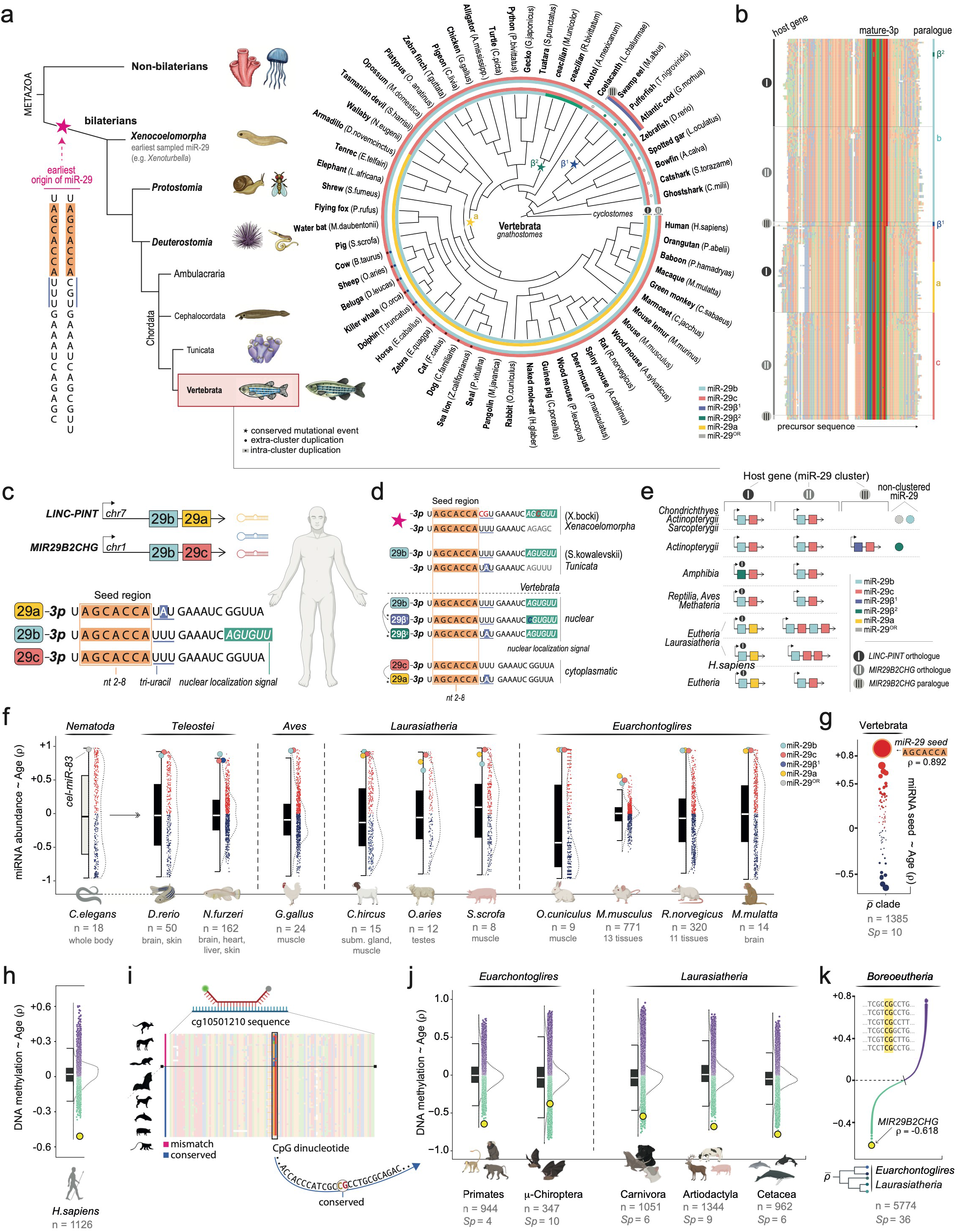
**The miR-29 age association extends across tissues and species**. **a**, Phylogenetic reconstruction of miR-29 loci, highlighting sequence changes and duplications. **b**, Illustrative precursor alignment with mature sequences and loci. **c**, Human chromosome 7 miR-29a/b1 and chromosome 1 miR-29b2/c loci, their mature products and shared AGCACCA seed; motifs are sequence annotations. **d**, Mature-sequence diversification outside the seed; β1/β2 denote comparative variants, not human *MIR29B1/B2* copies. **e**, Comparative cluster organization; empty positions denote unannotated copies. **f**, miRNA–age Spearman distributions across 1,403 profiles from 11 nonhuman species, highlighting miR-29; *C. elegans* miR-83 is a seed-related comparison. Sample sizes and tissues are indicated. **g**, Species- and group-balanced ranking across 1,385 profiles from ten nonhuman vertebrate species: AGCACCA ranks first among 76 seeds shared across five groups. **h**, Human age correlations for 37,554 mammalian-array probes, highlighting the most negatively associated probe, *MIR29B2CHG*-associated cg10501210 (1,075 contributing profiles, 13 tissues). **i**, Alignment of the interrogated region and target CpG across 60 species. **j**, cg10501210 in five group-specific CpG–age distributions; primates exclude human. **k**, Equal-group Boreoeutheria aggregate including human (5,617 contributing profiles, 36 species); cg10501210 remains the most negatively associated probe. Methylation correlations are averaged arithmetically across tissues within species and in Fisher-z space above species. Boxes show medians and interquartile ranges of feature coefficients; whiskers extend to 1.5 interquartile ranges. Coverage and contributing sample counts are distinguished in Supplementary Methods.

We compared age-associated miRNA rankings across 11 nonhuman species, including *C. elegans* miR-83 as a seed-related comparison (Fig. 2f, Supplementary Fig. 2b,c). Individual miR-29 family members showed recurring positive associations across vertebrate tissues and species, with variation in effect size. We then grouped miRNAs sharing the same seed across 1,385 profiles from ten nonhuman vertebrate species. Among 76 seeds represented in all five vertebrate groups, the conserved miR-29 seed had the strongest aggregate positive association with age (ρ = 0.892; Fig. 2g). Its association was positive in all ten species, and it retained first place after omission of any species or group and under alternative weighting schemes (Supplementary Methods). The leading position of miR-29 was therefore not driven by any single species and was established without the human discovery data.

DNA methylation provided a complementary view at the miR-29b2/c-associated locus. The probe cg10501210, annotated to *MIR29B2CHG* and included in a model for forensic age prediction in human blood ^11^, ranked as the most negatively age-associated site among 37,554 probes in humans (ρ = −0.509; Fig. 2h). Cross-species sequence comparisons assessed conservation of the CpG dinucleotide at the position corresponding to human cg10501210, restricting biological interpretation of methylation to species retaining this homologous CpG (Fig. 2i, Supplementary Fig. 2d–f). Among eligible mammals, this site again occupied the most negative aggregate age-association rank, with a coefficient of −0.618 (Fig. 2j,k). It remained first when human or all primates were excluded, first after every single-species omission, and first or second after omission of an entire analysis group (Supplementary Fig. 2g,h). Expression and locus-associated methylation therefore place miR-29 at an unusually prominent, evolutionarily recurrent position in molecular ageing.

## miR-29 inhibition opposes transcriptional ageing

We next asked whether miR-29 merely marks cellular ageing or can shift age-associated transcriptional programmes. Primary gingival fibroblasts from a 60-year-old donor received three rounds of pan-miR-29 inhibition or mimic treatment, each compared with its own scramble-control arm. Sampling at 72 and 96 hours after each round produced six timepoints (Fig. 3a). Independent experimental runs provided within-donor replication. Neither perturbation produced a detectable difference in the normalised growth trajectories over this interval (Fig. 3b). To test whether key responses extended beyond this preparation, we also examined independently sourced hTERT-immortalized gingival and SV40-immortalized pulmonary fibroblasts from other donors.

**Fig. 3.**
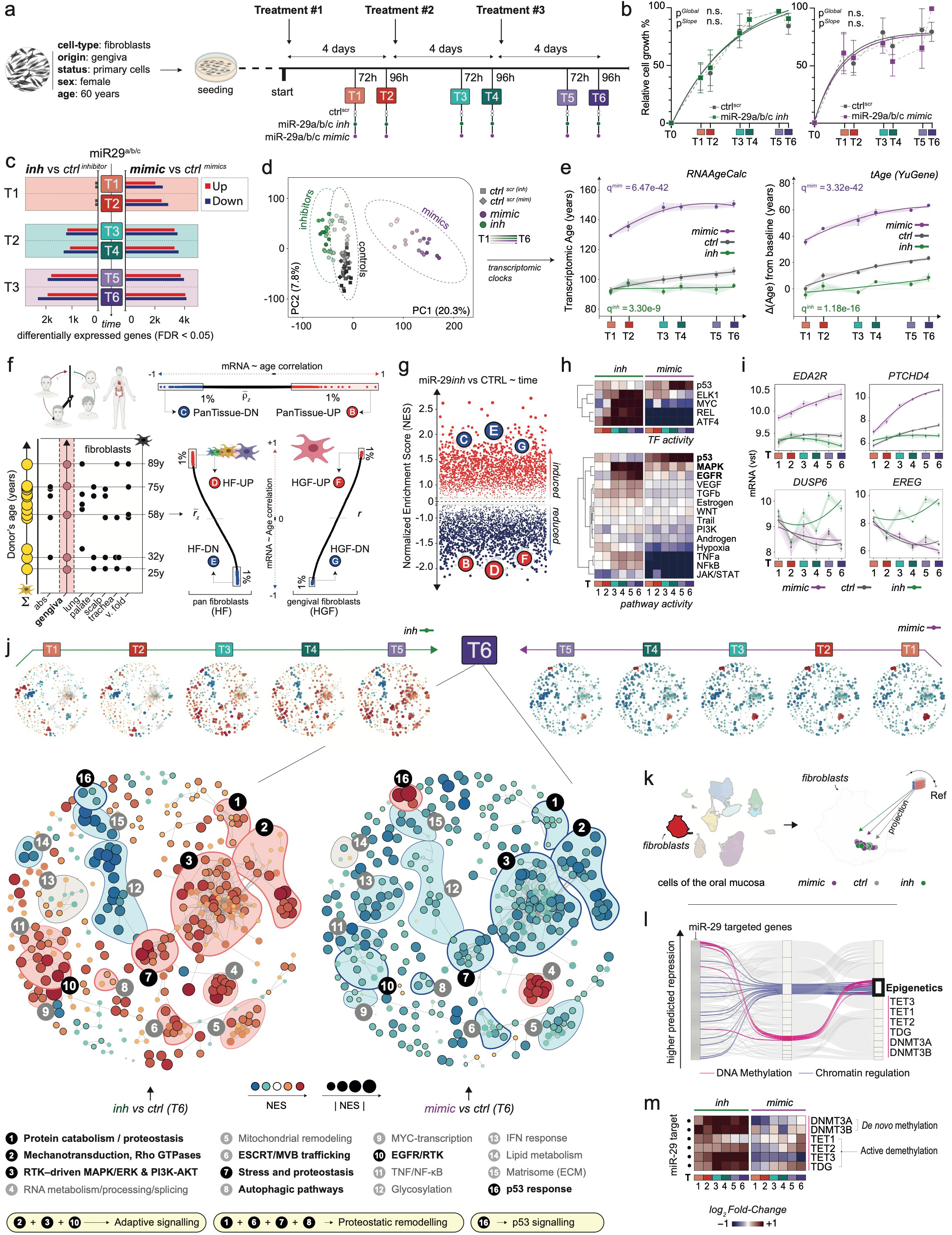
**miR-29 inhibition opposes transcriptional ageing programmes in primary fibroblasts**. **a**, Primary gingival fibroblasts from one 60-year-old female donor received three inhibitor or mimic treatments with matched scramble controls. T1–T6: 72, 96, 168, 192, 264 and 288 h. RNA-seq: five inhibitor/control and three mimic/control experiments; 96 samples. **b**, Growth: mean ± s.e.m., three experiments per arm (mimic, 264 h: n=2); exponential-plateau fits. **c**, Differentially expressed genes (batch-adjusted DESeq2; BH FDR <0.05). **d**, PCA of the 20,000 most variable batch-adjusted genes. **e**, RNAAgeCalc and tAge predictions; tAge is recentered on own-arm T1 controls. Global shifts use run-aware models versus pooled controls (five inhibitor, three mimic, eight control runs; four-test BH per clock). **f**, Independent age-positive/negative signatures: pan-tissue B/C, pan-fibroblast D/E, gingival-fibroblast F/G. **g**, Enrichment against inhibitor-by-time signed-Wald statistics: normalised enrichment scores (NES), multilevel probabilities and BH correction across 4,721 sets. **h**, VIPER transcription-factor and PROGENy pathway activities relative to own controls. **i**, p53- and EGFR/MAPK-associated expression; global DESeq2 tests versus matched controls, BH across genes. Controls are pooled for display. **j**, Enrichment networks: node colour indicates NES, size its magnitude and edges pathway overlap. **k**, Projection of all 96 transcriptomes onto an oral-mucosa single-cell reference. **l**, Functional classification of 149 annotated miR-29 targets. **m**, Treatment-versus-control log2 fold changes for six methylation/demethylation regulators. Points, bars and ribbons in e and i show mean ± s.e.m.; quadratic fits are descriptive. Ordinal spacing groups treatment cycles, not uniformly elapsed time. Matched-control clocks: Supplementary Fig. 3b; statistical procedures: Supplementary Methods.

The transcriptional responses were temporally asymmetric. Mimic treatment generated an early response, whereas the inhibitor response developed more gradually with repeated exposure (Fig. 3c and Supplementary Fig. 3a). The two perturbations also moved samples along distinct transcriptomic trajectories (Fig. 3d). The transcriptomic clocks RNAAgeCalc and tAge, which estimate age from gene-expression profiles, gave younger predictions under inhibition and older predictions after mimic treatment relative to their controls (Fig. 3e) ^12,13^. Separate control-arm displays and PASTA supported this contrast in predicted molecular age (Supplementary Fig. 3b) ^14^.

To test whether the result extended beyond trained predictors, we independently defined age-positive and age-negative programmes from normal human tissues, fibroblasts across seven anatomical sites, and gingival fibroblasts (Fig. 3f and Supplementary Fig. 3c,d). In the inhibitor-by-time response, all three age-positive programmes were enriched towards repression and all three age-negative programmes towards induction (all six BH q ≤ 2.52 × 10⁻⁴; Fig. 3g, Supplementary Fig. 3e). Their limited overlap showed that the agreement was not simply repeated testing of nearly identical gene lists. Rebuilding the pan-fibroblast programmes after tissue omission preserved the overall pattern (Supplementary Fig. 3f,g). Inhibition therefore opposed independently defined transcriptional ageing directions at both broad tissue and fibroblast-specific levels.

The transcriptional response also involved coordinated pathway and transcription-factor programmes. Expression-derived activities implicated EGFR/MAPK–ELK1 signalling and adaptive MYC- and ATF4-associated responses under inhibition. Mimic treatment instead elicited a prominent p53-associated response (Fig. 3h,i and Supplementary Fig. 3h). Time-resolved enrichment maps connected these changes to protein turnover, trafficking, stress responses and autophagy (Fig. 3j). Projection onto an oral single-cell reference retained the samples within the fibroblast compartment, indicating that these responses occurred without an overt shift in fibroblast transcriptional identity (Fig. 3k and Supplementary Fig. 3i).

Competitive testing of 285 annotated miR-29 targets detected a shift towards induction under inhibition and repression under mimic treatment from the first timepoint, before the broader inhibitor response emerged (Supplementary Fig. 3j). Among the established direct targets of miR-29 are enzymes that both add and remove DNA methylation, including DNMT3A/B, TET enzymes and TDG ^15,16^. Their coordinated transcriptional response to inhibitor and mimic treatment prompted us to examine whether these changes were accompanied by broader methylome remodelling (Fig. 3l,m).

## miR-29 constrains the amplitude of methylome change

Methylation profiles diverged most clearly from controls after repeated miR-29 inhibition (Fig. 4a). Within-condition temporal analyses identified 160,996 significantly changing assay probes under inhibition, compared with 23,072 in controls and none under mimic treatment at the retained threshold (Fig. 4b,c). The larger count under inhibition included both hypermethylating and hypomethylating probes.

**Fig. 4.**
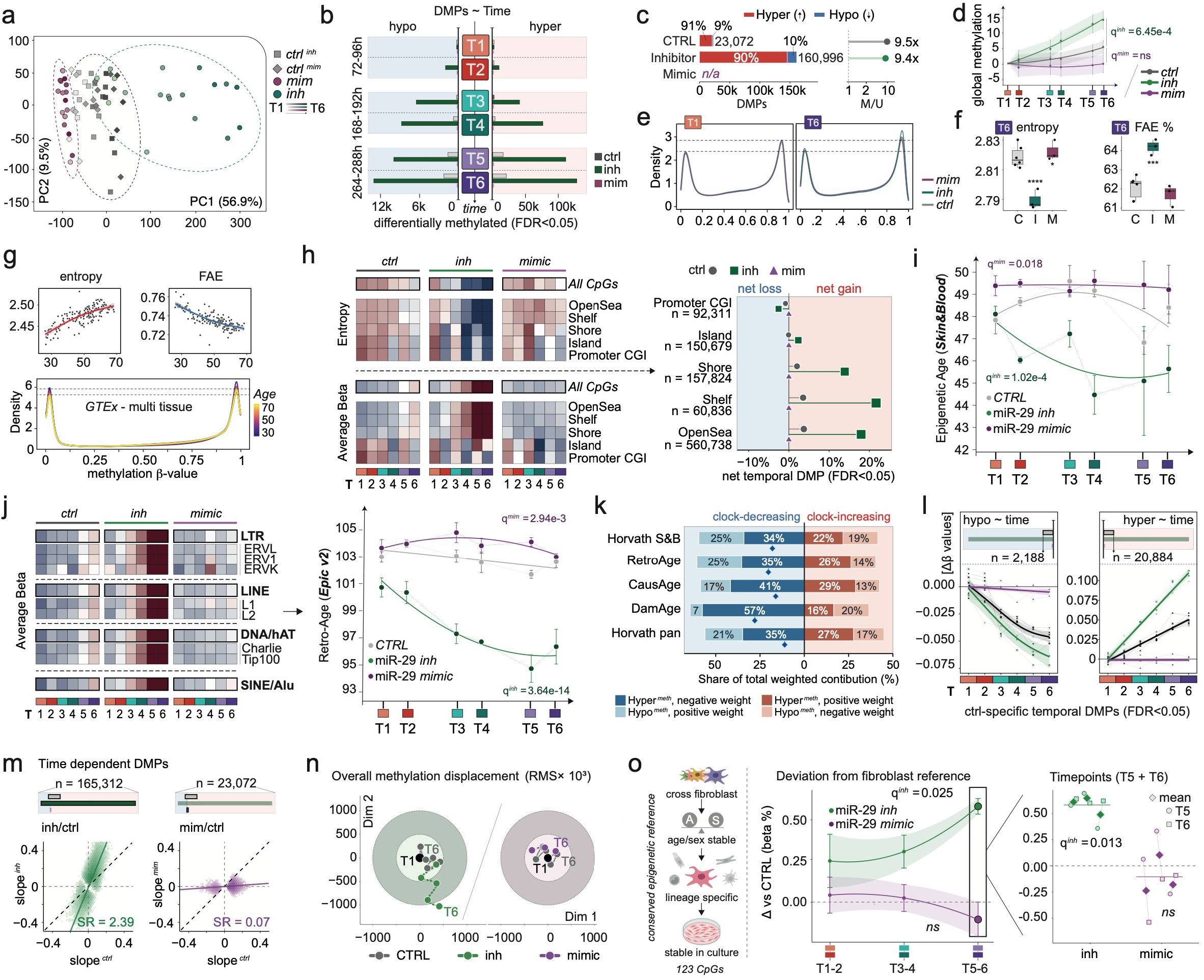
Lower epigenetic-clock scores accompany broader, asymmetric methylome remodelling. Three independent experiments per condition, six timepoints (72 primary-HGF methylomes). **a**, PCA of adjusted M values. **b**, Timepoint counts of temporal differentially methylated probes (DMPs), relative to CTRL T1. **c**, Temporal DMP counts and hyper/hypomethylation ratios (limma, BH FDR <0.05). **d**, Methylation changes from own T1 (1,000 × Δβ); 90% confidence intervals. **e**, T1/T6 methylation densities; bands indicate between-sample s.d. **f**, T6 entropy and fraction at extremes (FAE; β <0.2 or >0.8); three samples/treatment, six pooled controls. Mixed-model t contrasts; four-test BH. Boxes show medians/interquartile ranges; whiskers extend to 1.5 interquartile ranges. **g**, GTEx entropy/FAE versus DNAmAge and age-stratified methylation densities; 161 overlapping 15-sample windows (descriptive). **h**, Row-standardised compartment entropy/methylation. i,j, Net temporal DMP burden and repeat methylation, with Skin & Blood and RetroAge clocks, respectively. **k**, Time-integrated inhibitor-minus-control clock contributions by coefficient sign and methylation direction; percentages of absolute contribution, diamonds indicate net effects. **l**, Control-defined trajectories; GAM fits with 90% intervals. **m**, Temporal slopes; ratios from different selected sets are not directly comparable. **n**, RMS methylation displacement. **o**, 123-CpG fibroblast reference, treatment-minus-matched-control reference distances (β percentage points), cycle means ± s.e.m., late observations and run means (diamonds). Two-sided one-sample t tests assess full-series/T5–T6 means across three runs; two-arm BH per comparison. Clocks: mean ± s.e.m.; run-aware global shifts versus pooled controls, two-arm BH each. Interaction tests: pooled controls (d), four-test BH (l). Stars denote BH q <0.05 (*), <0.01 (**), <0.001 (***), <0.0001 (****); ns, q≥0.05.

The net effect included rising global methylation under inhibition, but the distributional response was not a uniform upward shift (Fig. 4d,e and Supplementary Fig. 4a). CpG methylation levels (β values) became more concentrated near the extremes, with fewer probes at intermediate values. This redistribution was accompanied by lower distributional entropy and a higher fraction of probes with low (<20%) or high (>80%) methylation (Fig. 4f and Supplementary Fig. 4b). Together, these changes opposed the ageing-associated patterns in normal tissues, shifting these distributional features towards a younger configuration (Fig. 4g).

The methylation response to miR-29 inhibition varied across genomic regions (Fig. 4h). Open-sea, shore and shelf regions showed a net excess of hypermethylating probes, whereas promoter CpG islands showed a net excess of hypomethylating probes (Fig. 4i, left). We assessed epigenetic age using the Skin & Blood clock, developed to improve age prediction in fibroblasts and other cell types, including cultured cells. Its predictions were lower under inhibition (Fig. 4i, right, and Supplementary Fig. 4c) ^17^. CausAge and DamAge also decreased, whereas AdaptAge increased (Supplementary Fig. 4d). AdaptAge captures protective age-associated methylation changes and rises during reprogramming; its increase under inhibition was consistent with the response reported in that setting ^18^. We next examined repeat-associated methylation, which contributes to transposable-element repression ^19^. Under miR-29 inhibition, mean methylation increased across the displayed repeat classes, accompanied by lower predictions from RetroAge, a clock built from retroelement-associated CpGs (Fig. 4j and Supplementary Fig. 4e) ^20^.

A lower clock estimate need not reflect a uniform shift towards younger predictions. We separated the methylation changes that lowered each clock’s prediction from those that raised it, accounting for each site’s contribution across the time course (Fig. 4k). All five decreasing clocks examined—Skin & Blood, RetroAge, CausAge, DamAge and Horvath pan-tissue—contained components acting in opposing directions. Clock-increasing components accounted for 36–45% of the total absolute weighted contribution. The younger prediction therefore reflected the balance between opposing components, not a uniform shift towards younger values. This decomposition describes the clock readout, not a percentage of biological rejuvenation.

We next asked whether inhibition changed the amplitude of methylome trajectories already present in control cultures. Probes changing with time in those cultures provided a common reference. Inhibition amplified the hypermethylating trajectory, increasing the change at the final timepoint approximately 2.3-fold relative to its own control, whereas mimic treatment restricted it (interaction q = 7.7 × 10⁻⁸ and 8.6 × 10⁻⁷, respectively; Fig. 4l). Mimic treatment also attenuated the hypomethylating trajectory. Across broader responsive-probe sets, temporal-slope comparisons were consistent with larger methylation gains and losses under inhibition (Fig. 4m and Supplementary Fig. 4f). Overall methylation displacement increased under inhibition and was constrained by mimic treatment (Fig. 4n). Together, these analyses show expanded remodelling in both directions, with the clearest evidence for amplification of pre-existing hypermethylating trajectories. miR-29 therefore controls the amplitude of methylome change rather than simply switching it towards methylation or demethylation.

## Greater plasticity alters lineage-associated methylation and stimulus responses

Such extensive remodelling raised a further question: does the methylome remain close to the patterns normally maintained in fibroblasts? We built a reference from independent fibroblast datasets, selecting 123 CpGs that distinguished fibroblasts from the other cell types tested and met stability criteria across age, sex, anatomical site and culture (Fig. 4o and Supplementary Methods). The reference included CpGs near *FAP*, *CCN1* and *DNM3OS*, genes implicated in fibroblast activation and tissue remodelling ^21,22,23^.

We measured each sample’s methylation distance from this reference relative to its matched control. Deviation increased across treatment cycles under inhibition, with a positive average difference from controls across the full series (two-arm BH q = 0.0253). At the final two timepoints, averaged within three independent runs, inhibition increased distance from the fibroblast reference by approximately 12% relative to matched controls (two-sided P = 0.00658; two-arm BH q = 0.0132; Fig. 4o). Mimic treatment did not produce a significant shift in either comparison. A uniform shift matching the global methylation change predicted a small decrease in reference distance, opposite to the observed increase (Supplementary Methods). These changes indicate increased divergence from the fibroblast methylation reference without an overt change in fibroblast transcriptional identity.

We tested whether this greater capacity for methylome change also enhanced the response to the GSK3 inhibitor CHIR99021. Three regions met the retained methylation-change threshold after CHIR99021 treatment, mapping to *HOXB3*/*HOXB-AS3*, *C1QTNF3*/*AMACR* and *DKK2* (Supplementary Fig. 4g) ^24^. Among these regions, *DKK2* combined strong statistical support with a direct role in WNT regulation. Matched profiles showed a stronger methylation response with combined miR-29 inhibition and CHIR99021 (Supplementary Fig. 4h). The combined response exceeded the sum of the two single-treatment changes in both the three-CpG core and the 16-CpG extended interval (q = 0.0096 and 0.0372 across the two region tests; Supplementary Fig. 4i). Inhibiting miR-29 therefore amplified the stimulus-induced methylation response at *DKK2*, extending its effect beyond trajectories already present in control cultures.

We next asked whether key responses extended to an independent donor background. In gingival fibroblasts from a different adult donor (hTERT), inhibition produced 3,632 temporal DMPs, predominantly hypermethylating. The inhibitor-defined hypermethylating and hypomethylating sets were enriched for changes in the same directions in control cultures, but not in mimic-treated cultures (Supplementary Fig. 4j). This pattern again supported enhanced remodelling under inhibition and constraint under mimic treatment. Transcriptomic clocks also reproduced the opposing responses: younger predictions under inhibition and older predictions under mimic treatment (Supplementary Fig. 4k). Applying the same 123-CpG reference showed a concordant increase in deviation under inhibition at the final cycle (two-arm BH q = 0.0409; Supplementary Methods).

To assess whether induced changes persisted after treatment stopped, we followed RNA and DNA methylation after inhibitor withdrawal in independently sourced SV40-immortalized pulmonary fibroblasts (Supplementary Fig. 4l). The RNA response largely subsided, whereas a substantial methylation component remained. At the final withdrawal timepoint, the RNA projection returned approximately to baseline (−6% of its T6 amplitude), whereas the DNAm projection retained 63% of its T6 amplitude (Supplementary Fig. 4m,n). Applying the same external fibroblast reference to this descriptive series also showed increased deviation at all eight observations, including after withdrawal (Supplementary Methods). These observations suggest that an induced epigenetic state can outlast its accompanying transcriptional response.

## The same miR-29–remodelling axis marks epigenetic deviation in cancer

The perturbation experiments linked miR-29 inhibition to lower molecular-age estimates, broader methylome remodelling and greater deviation from a lineage-associated reference. Cancer, where miR-29 is frequently reduced and stable cellular states are disrupted, provides a complementary context in which to examine this relationship. We therefore asked whether low miR-29 similarly accompanied younger epigenetic age and greater deviation from normal-tissue methylation profiles. In 8,428 unique TCGA tumour patients across 30 cancers, miR-29a, miR-29b, miR-29c and their composite showed the four strongest positive associations with DNAmAge among 1,764 miRNA entries (Fig. 5a, Supplementary Fig. 5a–c) ^25^. The cross-cancer pan-miR-29 association was also positive for age acceleration (ρ = 0.231), the clock residual after adjustment for chronological age.

**Fig. 5.**
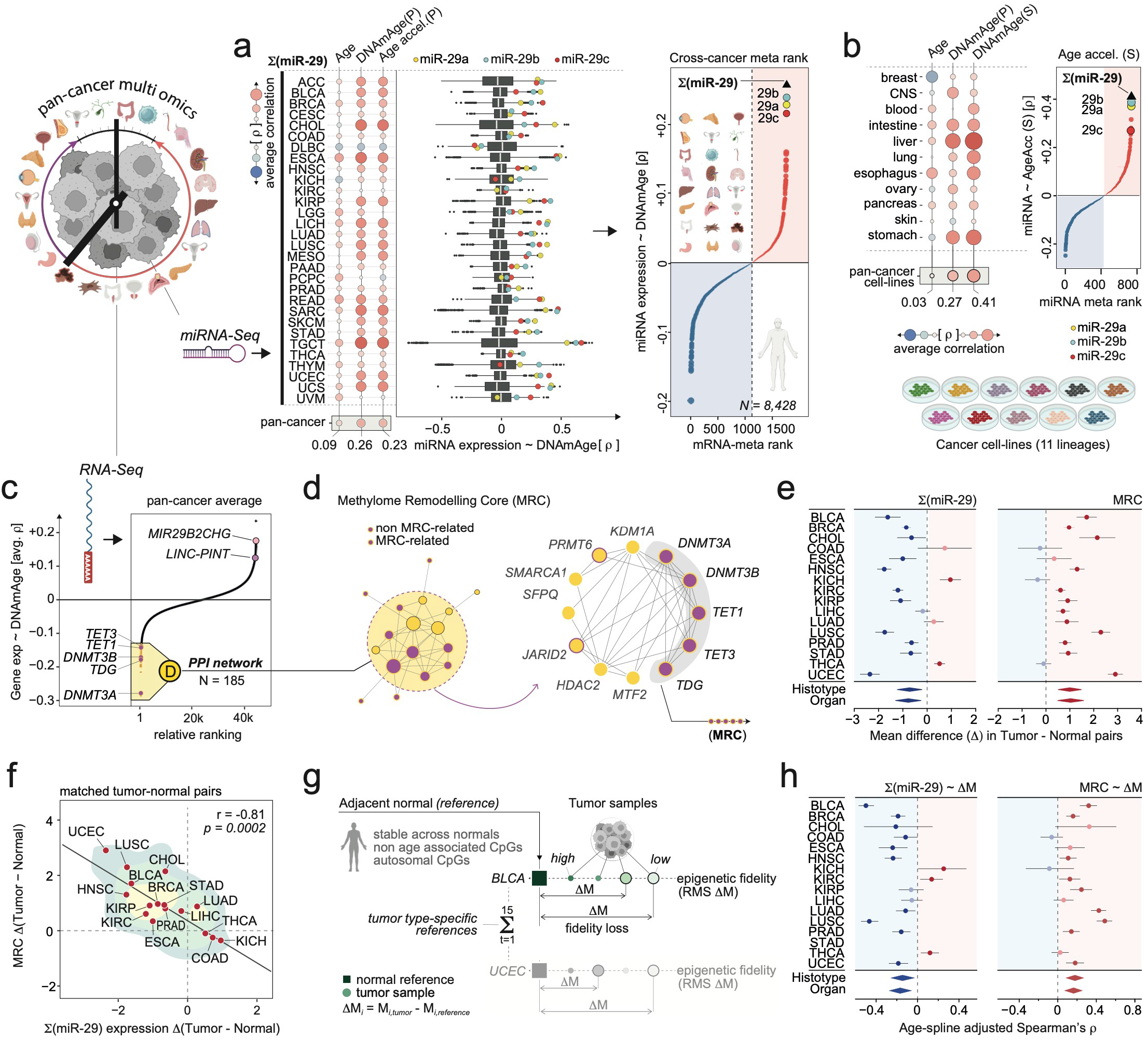
The miR-29–remodelling axis is associated with epigenetic fidelity in cancer. **a**, miRNA associations with chronological age, Horvath pan-tissue DNAmAge and acceleration in 8,428 tumour patients across 30 cancers; cancer-specific distributions and equal-cancer Fisher-z rankings highlight miR-29. **b**, Corresponding analysis of 429 CCLE lines across 11 lineages. Points show pan-miR-29 correlations with donor age and DNAmAge; ranking uses Skin & Blood acceleration. P, Horvath pan-tissue; S, Skin & Blood. **c**, Cross-cancer RNA–DNAmAge ranking; the negative three-standard-deviation tail contains 185 features. **d**, Interaction network; the methylome remodelling core (MRC) comprises DNMT3A, DNMT3B, TET1, TET3 and TDG. **e**, Mean standardised tumour-minus-normal pan-miR-29 and MRC expression differences in 580 matched patients across 16 histotypes. Bars show 95% t confidence intervals; diamonds show random-effects summaries across 16 histotypes or 13 organs. **f**, Association between 16 cancer-mean paired effects (Pearson r=−0.81, two-sided P=0.0002); contours show cancer-balanced patient-pair density, not confidence regions. **g**, Tissue-matched reference from stable autosomal CpGs in adjacent normals; deviation is the root-mean-square difference between tumour M values and normal-reference medians. **h**, miR-29/MRC associations with reference deviation across 15 histotypes after residualizing both variables on chronological-age splines. Bars show approximate 95% Fisher intervals; diamonds show REML/Knapp–Hartung summaries across histotypes or 12 organs. Forests use their respective available-assay cohorts. Boxes in a summarize miRNA coefficient distributions, not patient-level effect intervals. Ranking and forest pooling procedures are distinguished in Methods.

To examine this relationship outside the native tumour microenvironment, we analysed 429 CCLE cancer cell lines across 11 lineages (Fig. 5b, Supplementary Fig. 5d–f) ^26^. The pan-miR-29 composite ranked first among 735 miRNA entries for association with Skin & Blood age acceleration. Its recurrence in cultured cancer cells supports a tumour-cell-intrinsic component. For the DNAmAge associations in TCGA and CCLE, omission analyses showed that neither aggregate association was attributable to a single tumour type or cell lineage (Supplementary Fig. 5c,e).

A transcriptome-wide ranking against tumour DNAmAge identified 185 extreme negatively associated features (Fig. 5c, Supplementary Fig. 5g). Their interaction network contained multiple chromatin and methylation regulators (Fig. 5d). Five of the six enzymes examined in the fibroblast perturbations lay within this tail: *DNMT3A*, *DNMT3B*, *TET1*, *TET3* and *TDG*, all previously reported direct targets of miR-29 ^15,16^. We summarised the expression of these five genes in a methylome remodelling core (MRC) score, bringing together the methylation and demethylation arms implicated by the perturbation experiments.

To establish how this axis changes relative to the normal state, we compared strictly patient-matched tumour and normal expression profiles from 580 patients across 16 histotypes. The pooled comparisons revealed reduced miR-29 and increased MRC expression in tumours (Fig. 5e and Supplementary Fig. 5h). Across cancer-mean paired effects, larger miR-29 decreases accompanied larger MRC increases (Pearson r = −0.806; Fig. 5f), and the relationship persisted in leave-one-cancer-out analyses (Supplementary Fig. 5i). In most analysed histotypes, tumours therefore combined reduced expression of the proposed constraint with increased expression of the machinery for bidirectional methylome change.

Having linked reduced miR-29 to increased expression of its methylome-remodelling targets, we asked whether this axis also accompanied deviation from normal-tissue methylation profiles. For each of 15 eligible histotypes, we selected autosomal CpGs that met stability criteria in normal tissues and measured tumour deviation from the resulting reference (Fig. 5g and Supplementary Fig. 5j). These tissue-specific references measured how far each tumour’s methylation profile had diverged from patterns normally maintained in its tissue of origin.

After accounting for chronological age, methylation divergence from tissue-matched normal profiles correlated negatively with miR-29 expression (12 of 15 histotypes; pooled ρ = −0.145, 95% confidence interval −0.258 to −0.028) and positively with MRC scores (13 of 15 histotypes; pooled ρ = 0.179, 95% confidence interval 0.084 to 0.270; Fig. 5h). Effect sizes varied across cancers. The miR-29 association remained similar after additional adjustment for age acceleration (ρ = −0.150). Age acceleration itself showed a near-zero pooled association with reference deviation, with heterogeneous effects across cancers (Supplementary Fig. 5k). A clock residual and reference fidelity therefore capture different aspects of the methylation state. Together, the tumour–normal shifts and reference-deviation associations connect both sides of the miR-29–remodelling axis to epigenetic disruption across human cancers.

## Discussion

Intersecting transcriptional programmes of physiological ageing, induced reprogramming and early development led us to miR-29, whose exceptional association with age recurs across tissues and species. miR-29 directly targets enzymes that both add and remove DNA methylation ^15,16^. Our findings place this regulatory role at the intersection of molecular age, epigenetic plasticity and fidelity. Releasing miR-29 restraint expanded methylome remodelling, opposed transcriptional ageing programmes and lowered several molecular-age predictions. Lower epigenetic-clock estimates emerged from the unequal balance of opposing methylation changes, rather than a uniform reversal of the ageing methylome. These younger predictions coexisted with greater deviation from lineage-associated methylation patterns, without an overt shift in fibroblast transcriptional identity. The distinction between the capacity to change and the state reached by that change is therefore central: molecular age and fidelity to a tissue-associated reference capture distinct outcomes when assessing rejuvenation.

Earlier studies linked miR-29 to age-dependent DNA-damage responses, cross-tissue ageing, methylation enzymes and reprogramming ^15,16,27,28,29^. Neural studies identified constraints on adult neuronal plasticity and memory-related methylation ^30,31^. In earlier fibroblast work, depletion favoured demethylation and brought reprogrammed cells closer to an embryonic-stem-cell methylation reference ^16^. Here, inhibition instead produced a net methylation gain and increased distance from a stable somatic reference. These comparisons address different epigenetic destinations. Our study brings these previously separate observations into a common framework, positioning miR-29 not simply as another age-associated factor, but as a central point of convergence between ageing and rejuvenation. Its exceptional age association across tissues, species and molecular layers is linked by experimental perturbation to functional effects on methylome remodelling, molecular age and the preservation of epigenetic fidelity.

Excess miR-29 can itself be detrimental: its overexpression induces ageing-like phenotypes and early mortality in mice ^32^. Conversely, loss of miR-29 control over DNMT3A during brain maturation increases non-CG methylation and disrupts neuronal function ^33^. These findings indicate that neither increasing nor suppressing miR-29 is uniformly beneficial.

These findings support a model in which rising miR-29 helps stabilise established epigenetic states at the expense of youthful plasticity. Age-associated demethylation at the miR-29b2/c locus raises the possibility that epigenetic change itself contributes to miR-29 induction. Such coupling engages a feedback response that restrains methylome remodelling to protect against further epigenetic drift, linking the accumulation of epigenetic change to the progressive restriction of plasticity. This stabilising response becomes maladaptive when it limits the remodelling needed for methylome maintenance and adaptation to new stimuli. miR-29 contributes to the aged cellular state by promoting age-associated transcriptional programmes while reinforcing epigenetic rigidity. miR-29 inhibition increased methylation at repeat-associated CpGs (Fig. 4j), contrasting with the methylation loss reported at transposable elements during ageing ^34^. Releasing this constraint can reopen access to younger molecular configurations, but does not ensure a return to a faithful epigenetic state. This interpretation is consistent with adaptive miR-29 induction limiting neuronal iron accumulation in killifish ^35^, protective effects in other contexts and broader proposals connecting ageing to regulatory constraint ^36,37^.

Independently sourced gingival fibroblasts from a different adult donor (hTERT) showed the same combination of younger transcriptomic-age predictions, increased methylome remodelling and greater deviation from the fibroblast reference under miR-29 inhibition. A separately sourced pulmonary fibroblast model further showed persistent methylation changes and increased reference deviation after withdrawal. Reference deviation measures methylation distance, not a functional loss of identity, and the withdrawal series provides a descriptive view of persistence.

The recurrence of the miR-29–epigenetic age association in cultured cancer cells, outside the native tumour microenvironment, supports a tumour-cell-intrinsic component. The cancer analyses extend the relationship with reference-defined fidelity beyond the fibroblast signature. Pooled tumour– normal comparisons show reduced miR-29 and increased remodelling-enzyme expression. Across 15 histotypes, lower miR-29 and higher remodelling-enzyme expression accompany greater deviation from tissue-matched normal methylation references. These complementary findings connect the same regulatory axis to epigenetic disruption across human malignancies; its causal role in tumour development remains to be tested.

By amplifying both ongoing remodelling and stimulus-induced methylation changes, miR-29 inhibition raises the possibility of opening a transient window of plasticity. Such a window could enhance responses to anti-ageing and rejuvenating interventions by overcoming epigenetic resistance to change. The opportunity is to harness this trade-off by reopening the capacity for change while keeping epigenetic drift in check.

## Methods

### Study design and experimental models

The study combined public-data analyses with miR-29 perturbation in human fibroblasts. The primary human gingival fibroblasts (HGF; ATCC PCS-201-018) were derived from one 60-year-old female donor. Independent experimental runs used this same donor-derived culture. Two independently sourced models were male-derived hTERT-immortalized HGF (ABM T0026) ^38^ and SV40-immortalized pulmonary fibroblasts from a 65-year-old female donor (HPF; ABM T0490). The primary-HGF RNA-seq experiment included five inhibitor/control runs and three mimic/control runs, with six timepoints per condition (96 samples). The corresponding DNA-methylation experiment included three runs per arm at six timepoints (72 samples). Each run contributed one analysed sample per condition and timepoint, and each treatment arm had its own non-targeting control. Growth measurements used three experiments per arm, except mimic at 264 h (n = 2). T1–T6 corresponded to 72, 96, 168, 192, 264 and 288 h. Further model-specific sample structures are described below and in Supplementary Methods. No newly generated experimental samples were discarded; analysis-specific eligibility and feature-filtering criteria are described below and in Supplementary Methods.

### Statistical analysis

Statistical tests, model specifications and multiple-testing corrections are described with each analysis below and in Supplementary Methods. Tests were two-sided unless stated otherwise. Experimental runs were the units of replication in replicated perturbation experiments; repeated measurements were accounted for in the models or summarised within run.

### Cell culture and miR-29 perturbation

Primary HGF were maintained in Fibroblast Basal Medium (ATCC PCS-201-030) supplemented with the Fibroblast Growth Kit–Low Serum (ATCC PCS-201-041) and 100 U ml⁻¹ penicillin– streptomycin (ABM G255). Immortalized HGF and HPF were maintained in PriGrow III medium (ABM TM003), 10% fetal bovine serum (Euroclone) and penicillin–streptomycin on extracellular-matrix-coated plates (ABM G422). Cultures were maintained at 37 °C and 5% CO₂. All cultures were purchased directly from the manufacturers, authenticated and routinely tested for mycoplasma contamination, with negative results.

The three miR-29 family members were targeted together using synthetic miR-29a-3p, miR-29b-3p and miR-29c-3p inhibitors (MH12499, MH10103 and MH10518; Life Technologies) or mimics (MC12499, MC10103 and MC10518). Across all fibroblast models, inhibitor and mimic pools were used at final total oligonucleotide concentrations of 50 nM and 25 nM, respectively. Corresponding non-targeting controls were used at 50 nM for inhibitor experiments and 25 nM for mimic experiments. Non-targeting inhibitor (4464076) and mimic (4464058) reagents provided the corresponding controls. Transfection used Lipofectamine RNAiMAX (13778150) in Opti-MEM I Reduced Serum Medium with GlutaMAX (51985034), according to the manufacturer’s instructions.

### RNA sequencing and preprocessing

RNA was extracted using the miRNeasy Mini Kit (QIAGEN 217004). RNA concentration and quality were assessed using Qubit RNA High Sensitivity assays and an Agilent 4200 TapeStation. Libraries were prepared using the TruSeq Stranded Total RNA protocol and sequenced on an Illumina NextSeq 500. Samples from different treatment groups were distributed across sequencing runs to avoid a one-to-one correspondence between treatment and sequencing batch. Single-end reads were aligned to the GRCh38 primary assembly with STAR v2.7.9a ^39^, using GENCODE v45 primary-assembly annotation and two-pass mode (--twopassMode Basic). Gene counts were generated with -- quantMode GeneCounts; downstream analyses used the reverse-stranded counts in column 4 of ReadsPerGene.out.tab.

### DNA methylation profiling

DNA was extracted using the QIAamp DNA Mini kit (QIAGEN 51104), quantified with Qubit dsDNA High Sensitivity assays, bisulfite-converted with the EZ DNA Methylation Kit (Zymo Research D5001), and profiled on Infinium MethylationEPIC v2 arrays using an Illumina iScan. Sample positions on the arrays were randomised.

### Convergence of ageing and rejuvenation programmes

Reprogramming RNA-seq counts from GSE165176 were analysed using DESeq2 ^40^. The comparison comprised 36 samples from six donors, including two neonatal donors, at days 11, 13 and 15; the separate methylation trajectory contained 22 records from four adult donors. A model containing group, donor and numeric time estimated the SSEA4-positive versus CD13-positive branch main effect over this interval, without a group-by-time interaction. Reprogramming-down genes were those below the mean Wald statistic minus three standard deviations (531 genes).

Developmental RNA-seq from GSE101571, GSE157834, GSE71318, GSE65481 and GSE133854 was variance-stabilized, and cohort effects were removed while preserving developmental stage. Twenty principal components, four k-means clusters and Slingshot defined a developmental pseudotime ^41^. The trajectory ordered RNA profiles toward increasing developmental stage. Spearman correlations between expression and pseudotime were Fisher-transformed; the lower three-standard-deviation tail defined the 175-gene developmental programme.

Normal-tissue age associations used precomputed GTEx-derived gene–age correlations. Tissue-level analyses included adjustment for sex and anatomical sublocation and age-constrained resampling. Gene correlations were combined on the Fisher-z scale; the upper three-standard-deviation tail defined the 179-gene ageing programme. Aggregation and resampling procedures are detailed in

Supplementary Methods. The chronological-age and DNAmAge rank comparison used the same seven tissue categories, with endpoint-specific eligible samples; it did not require identical biospecimens across endpoints. The separate *LINC-PINT* association analysis used strictly matched RNA–methylation samples. Healthy-tissue mature-miRNA analyses used the processed GTEx v10 small-RNA release (17 March 2025; dbGaP phs000424), without reprocessing its raw reads. Mature-miRNA developmental and reprogramming analyses used the published expression tables and their stage or time assignments.

### Comparative miRNA and methylation analyses

Public small-RNA data were analysed from raw reads reprocessed with nf-core/smrnaseq v2.4.1 (https://nf-co.re/smrnaseq/2.4.1) ^42^ or from published expression matrices, as specified in Supplementary Methods. Reprocessing used species-specific mature and hairpin sequences from miRBase ^43^, with the NFINgb genome and miRNA annotation used for Nothobranchius furzeri ^44^. Adapter trimming and read-length processing used fastp with dataset-specific settings ^45^.

Species-specific miRNA expression was related to chronological age using Spearman correlation. Mature sequences were mapped to seeds, collapsed in Fisher-z space within species, and combined with equal species weight within five analysis groups and equal group weight in the final summary. Seeds required representation in at least one species of each group, retaining 76 of 1,966 observed seeds. The seed analysis excluded human and comprised 1,385 profiles from ten species. Sequence and genomic-locus comparisons used accession-level inventories containing cluster membership, precursor and mature sequences, assembly accessions and strand-specific genomic coordinates. The vertebrate topology was obtained from Open Tree and retained as a resolved tree. The cg10501210 target-region alignment was generated with MAFFT and BMGE through the NGPhylogeny.fr PhyML-SMS/OneClick workflow ^46^.

Pan-mammalian methylation analyses used normalised Mammalian Methylation Array data from GSE223748. Samples required a recorded age and age-confidence scores above 80, and species– tissue strata required at least ten samples. Species required at least one eligible tissue. For each of 37,554 probes, Spearman correlations with age were calculated within strata. Tissue coefficients were averaged arithmetically within species; species or balanced genus-level units were then combined in Fisher-z space within five analysis groups, each weighted equally: Primates, Chiroptera, Carnivora, non-cetacean Artiodactyla and Cetacea. Leave-one-species-out and leave-one-group-out analyses repeated the aggregation and ranking over the full probe universe. The primate display excluded human, whereas the full aggregate included human. Target conservation was assessable in 66 species: 41 retained the CpG and 25 were classified as non-conserved, including 19 with a disrupted dinucleotide and six without an identifiable homologous probe-target region in the available genome. Eleven species lacking a suitable exact-species reference were excluded. Conserved versus non-conserved comparisons were treated as assay-context controls, not as measurements of methylation loss at an identical conserved site.

### Primary-HGF transcriptomic analyses

Timepoint-specific differential expression used raw counts in separate DESeq2 models for inhibitor and mimic arms, including treatment-by-timepoint groups and experimental run. Contrasts compared treatment with its own control at the same timepoint. Differential expression required BH-adjusted P < 0.05; no fold-change cutoff was imposed. The independent-filtering setting supplied to DESeq2 results was alpha = 0.001. Global expression comparisons were fitted separately for the inhibitor and mimic arms, including treatment group, categorical timepoint and experimental run as additive terms. Treatment-versus-own-control Wald tests retained transcriptome-wide BH adjustment and independent filtering. Visualisation used variance-stabilized expression with design-protected adjustment for experimental arm and run. PCA used the 20,000 most variable genes, centred and scaled.

Age-associated fibroblast signatures were defined independently of the perturbation data using 34 samples from 15 individuals and seven anatomical sites in GSE140523 ^47^. Within each tissue, expression and age were separately regressed on sex with an intercept, and Pearson correlations were calculated between residuals. Equal-weight Fisher-z aggregation required at least five finite tissue estimates. Strict upper and lower 1% tails, with positive and negative sign requirements, defined pan-fibroblast signatures; *LINC-PINT* and *MIR29B2CHG* were excluded without replacing genes. Gingival signatures used the same partial-correlation procedure in the five gingival samples. The pan-fibroblast and gingival signatures comprised 149/149 and 125/125 up/down genes. The pan-tissue sets comprised 354/355 unique genes.

Temporal enrichment used the signed Wald statistic for the inhibitor-by-time term. Normalized enrichment scores were obtained with fgseaSimple and 10,000 permutations; significance was estimated with fgseaMultilevel, followed by BH adjustment across the 4,721 tested sets ^48^. Transcription-factor and pathway activities were estimated using DoRothEA/VIPER and PROGENy, respectively ^49,50,51^. Activity heatmaps show mean treatment-minus-own-control scores at each timepoint. Bulk transcriptomes were projected onto a published oral single-cell reference (GSE164241); this analysis assessed broad transcriptional identity rather than identifying new cell populations.

### Molecular-age models

RNA ages were estimated using the published RNAAgeCalc, tAge and PASTA models with fixed parameters ^12,13,14^. tAge used the human-inclusive YuGene model and a per-gene negative-control reference calculated separately within each arm. Display transformations, including the 122.5 model-scale multiplier and T1-control recentering, were distinguished from the prediction input and statistical tests. Skin & Blood, Horvath pan-tissue, RetroAge, CausAge, DamAge and AdaptAge methylation predictors used their published coefficients ^3,17,18,20^. CausAge incorporates CpGs prioritised for causal effects on ageing-related traits using Mendelian randomization, whereas DamAge and AdaptAge separate methylation changes classified as damaging and adaptive, respectively. Skin & Blood and causal-clock inputs used adjusted, base-CpG-collapsed beta values; RetroAge retained EPIC v2 assay identifiers. Missing features in the causal clocks were omitted without rescaling the remaining coefficients, and coverage is reported in Supplementary Methods. Clock curves report global average-shift q values, using pooled controls in the main figures and matched controls in control-resolved supplementary panels; both analyses retain run-aware models and their stated correction families (Supplementary Methods).

### Genome-wide methylation analysis

IDAT files were processed with the openSesame function in SeSAMe using the CDB preprocessing sequence ^52^. Beta values and M values, M = log₂[beta/(1−beta)], were retained at assay-probe and collapsed-CpG resolution. Replicate-probe collapsing was performed separately on the beta and M scales. Temporal DMP analyses used noncollapsed assay-level M values, whereas raw collapsed beta values were used for global methylation and the fibroblast-reference score.

Temporal methylation models used limma with numeric ordinal time, condition, their interaction, and experimental arm, accounting for experimental-run blocks through duplicateCorrelation ^53^. Empirical-Bayes contrasts estimated control, inhibitor and mimic slopes; the control slope used 36 samples from both control arms, whereas each treatment slope used 18 samples. BH-adjusted P < 0.05 defined temporal DMPs, with direction determined by the slope sign. For timepoint bars, BH correction was recalculated within each condition’s selected temporal-DMP set; these bars therefore describe a conditional testing family. Through-origin slope comparisons used the union of inhibitor- and control-significant probes for inhibition and control-significant probes for mimic, which had no significant temporal probes.

Global methylation was the sample mean across complete collapsed CpGs, analysed with run-aware mixed models and contrasted with each condition’s T1 baseline. The displayed scale is 1,000 times the change in mean beta. Methylation entropy was calculated over twenty equal-width beta bins, and the fraction at extremes was the fraction with beta < 0.2 or beta > 0.8. These are sample-level distributional summaries and do not quantify cell-to-cell heterogeneity. Genomic-compartment and repeat summaries used genomic annotations and adjusted noncollapsed beta values. Their heatmaps were standardised within rows. The methylation-displacement representation used temporal centroid differences across the 50,000 most variable finite M-value features, preserving step RMS magnitude in a common two-dimensional representation.

### Clock-component decomposition

For each clock CpG and experimental run, inhibitor-minus-matched-control beta differences were integrated by signed trapezoids over 72–288 h and divided by the 216-h duration. The time-averaged difference was multiplied by the published clock coefficient. Contributions were assigned according to methylation-change direction and coefficient sign, yielding two predictor-decreasing and two predictor-increasing routes. Mean route contributions across runs were divided by the mean total absolute CpG contribution. For models with nonlinear age transformations, decomposition preceded that transformation. Percentages represent contributions to the additive clock predictor.

### External fibroblast methylation reference

Reference features were derived from external fibroblast datasets. From GSE77135, autosomal, platform-compatible sites required at least 90% detection, an absolute beta-value difference ≤0.05 between scalp-derived and dura-mater-derived fibroblasts, absolute age slope ≤0.001 beta per year and absolute sex contrast ≤0.05 beta. Of 384,494 screened sites, 222,344 met these conditions. GSE179847 normal/control fibroblast cultures retained 169,861 sites with at least 90% detection and predicted full-range culture changes within ±0.05 beta in both unadjusted and batch-adjusted fits. Cell-type association was evaluated in GSE31848 using a balanced fibroblast-versus-epithelial/smooth-muscle contrast, sex-adjusted BH q ≤0.20, concordant adjusted and unadjusted directions, and absolute differences ≥0.10 beta in both fits. The same direction and magnitude were required against purified immune-cell profiles from GSE110554. The resulting reference contained 123 sites: 36 fibroblast-high and 87 fibroblast-low.

For each sample, reference deviation was D = mean_j_|β_j_−r_j_|, where r_j_ was the external fibroblast median for CpG j. Matched effects were 100 × (D_treatment_−D_control_) within run, timepoint and arm. Displayed cycle means first averaged T1/T2, T3/T4 or T5/T6 within run. For inference, matched effects were averaged within each run over all six timepoints or over T5/T6, yielding three run means per arm for each comparison. Two-sided one-sample t tests assessed mean displacement from the matched-control null, with BH adjustment across inhibitor and mimic separately for each temporal summary. The full-series test assessed average displacement, not change with time. The between-arm comparison used Welch’s test. This beta-distance score differs from the M-value RMS cancer score described below.

### CHIR99021, hTERT and withdrawal experiments

The CHIR99021 methylation experiment compared non-targeting control, miR-29 inhibitor, control plus CHIR99021 and inhibitor plus CHIR99021. CHIR99021 (10 μM) was added at the medium changes following the second and third transfection rounds, for a total of two additions. No CHIR99021 was added after the first round to allow time for downstream targets to respond to miR-29 inhibition. DMRcate discovery used all available samples with group and biological experiment in the design ^24^. CpG annotation used FDR 0.10, followed by regional smoothing with lambda = 1,000 and C = 2. Three of four CHIR99021-versus-control regions met the absolute mean delta-beta threshold of 0.10. Matched regional tests used the three complete experiments. Within each *DKK2* interval, beta was averaged across CpGs per sample. The factorial contrast was 100 × [(INH+CHIR99021−CHIR99021)−(INH−CTRL)]. One-sample t tests across three experiments and BH correction across the nested three- and sixteen-CpG intervals assessed beta-scale nonadditivity conditional on locus selection. The regional readout measures DNA methylation, not WNT transcriptional activity. hTERT HGF analyses comprised a separate model. RNA clocks used 42 samples from four inhibitor/control and three mimic/control runs at three timepoints. Inhibitor-defined temporal methylation sets were assessed competitively against control and mimic moderated t statistics using cameraPR with use.ranks = FALSE and inter-probe correlation 0.01 ^54^; neither control nor mimic yielded a significant temporal probe set. Reference-deviation analysis used 36 methylomes from three runs per arm, with treatment and matched control collected 96 h after each of three cycles. The same 123 CpGs and external centroid were applied without refitting. Final-cycle and full-series mean effects were tested across three run-level matched differences per arm, with two-sided t tests and separate two-arm BH corrections. The HPF series comprised one inhibitor and one control sample at each of eight observations: six during repeated treatment and two after withdrawal. RNA and DNAm retention projected each observed effect vector onto the observed T6 vector. The DNAm core was selected in a model that included post-withdrawal observations. The envelopes represent sensitivity across seven nested feature sets for RNA and five for DNAm, not biological confidence intervals.

### Cancer and cell-line analyses

TCGA tumour-age analyses retained one sample per patient, preferring primary over metastatic tumour, and required miRNA, age and Horvath DNAmAge. Adjacent normals were excluded from this cohort. The resulting cohort contained 8,428 patients in thirty cancers. Within each cancer, correlations from 100 random 80% subsamples were averaged in Fisher-z space, followed by equal-cancer aggregation. RNA associations were analysed similarly; 185 of 44,398 Ensembl features fell below the mean correlation minus three standard deviations. The methylome remodelling core (MRC) score comprised *DNMT3A*, *DNMT3B*, *TET1*, *TET3* and *TDG*. These five of the six regulators examined in Fig. 3m lay within the negative tail.

The CCLE analysis included 429 lines in eleven lineages, requiring complete donor age, Horvath and Skin & Blood predictions and at least fifteen lines per lineage. Correlations were aggregated with equal lineage weight. Paired tumour–normal expression analyses separately required both RNA and miRNA in both tissues from the same patient, yielding 580 pairs across sixteen histotypes. For these comparisons, each MRC gene was standardised across the combined tumour–normal universe within cancer, the five gene scores were averaged and the composite was standardised again. Standardized tumour-minus-normal effects were tested within cancer; Pearson correlation summarised the association between the sixteen cancer-mean miR-29 and MRC effects.

Normal-reference methylation analyses included fifteen histotypes. Eligibility for an own-tissue reference required at least five strict tumour–adjacent-normal methylation pairs; STAD had two and was excluded. All available adjacent normals were used to construct each eligible reference. KICH, which lacked adjacent-normal methylomes, was the exception and used a shared kidney reference containing 160 KIRC and 45 KIRP normals. Autosomal CpGs required ≤5% missingness before imputation, normal-sample SD(beta) ≤0.05, |Pearson correlation with age| ≤0.25 and absolute beta change per decade ≤0.01. Beta values were clipped to [10⁻⁶,1−10⁻⁶] before M conversion. Tumour reference deviation was the root mean square of M-value differences from normal-sample median M values. Both variables in fidelity associations were residualised against a three-degree-of-freedom natural spline of age before Spearman correlation. Fisher-z estimates were pooled using random-effects REML with Knapp–Hartung inference; assay-specific overlap cohorts were retained separately.

## AI-assisted support

Large language model tools (ChatGPT/Codex, OpenAI; Claude, Anthropic) assisted manuscript drafting. The authors reviewed the outputs and retain responsibility for the analyses, interpretation and final text.

## Data availability

Newly generated RNA-sequencing and DNA-methylation data have been deposited in GEO and will be made publicly available. Processed analysis inputs are deposited in Zenodo and will also be made publicly available. Supplementary Figures 1-5 are appended to this manuscript. Supplementary Methods and supplementary references are provided separately. Complete analysis tables will be made publicly available. Accessions and source publications for reused public datasets are listed in Methods and Supplementary Methods. Individual GTEx donor-age metadata are excluded from the shared inputs and require authorised dbGaP access (phs000424).

Selected study results will also be explorable through Tabula Ages, an interactive companion resource that will be made publicly available.

## Code availability

Analysis code, figure-specific reproducibility documents and input instructions are deposited in a GitHub repository. The repository link will be provided with the peer-reviewed publication. Processed inputs are deposited separately in Zenodo (see Data availability).

## Ethics and consent

Primary human gingival fibroblasts were purchased from ATCC (PCS-201-018), and hTERT-immortalised gingival fibroblasts and SV40-immortalised pulmonary fibroblasts were purchased from Applied Biological Materials (T0026 and T0490, respectively). No participants were recruited, no human tissues were collected specifically for this study and no experiments on live animals were performed. Controlled-access GTEx donor-age metadata were accessed through dbGaP under approved project 44958, ‘Age-associated changes in gene expression across normal human tissues’, at the Institute of Oncology Research (IOR), Switzerland. Other analyses used previously published datasets.

## Author contributions

I.C. and M.C.B. performed the wet-laboratory experiments and contributed to the conceptual development of the study. G.A.C. contributed to the conceptual development of the study and manuscript drafting. A.C.S. developed the web-based framework for data analysis and visualisation.

A.S. contributed to network and cancer-data analyses. L.G. contributed to raw-data processing. A.V. contributed to phylogenetic analyses. M.B. conceived and designed the study, supervised the project, led and performed the computational analyses, prepared the figures and supplementary materials, and wrote the manuscript.

---

**Supplementary Fig. 1.**
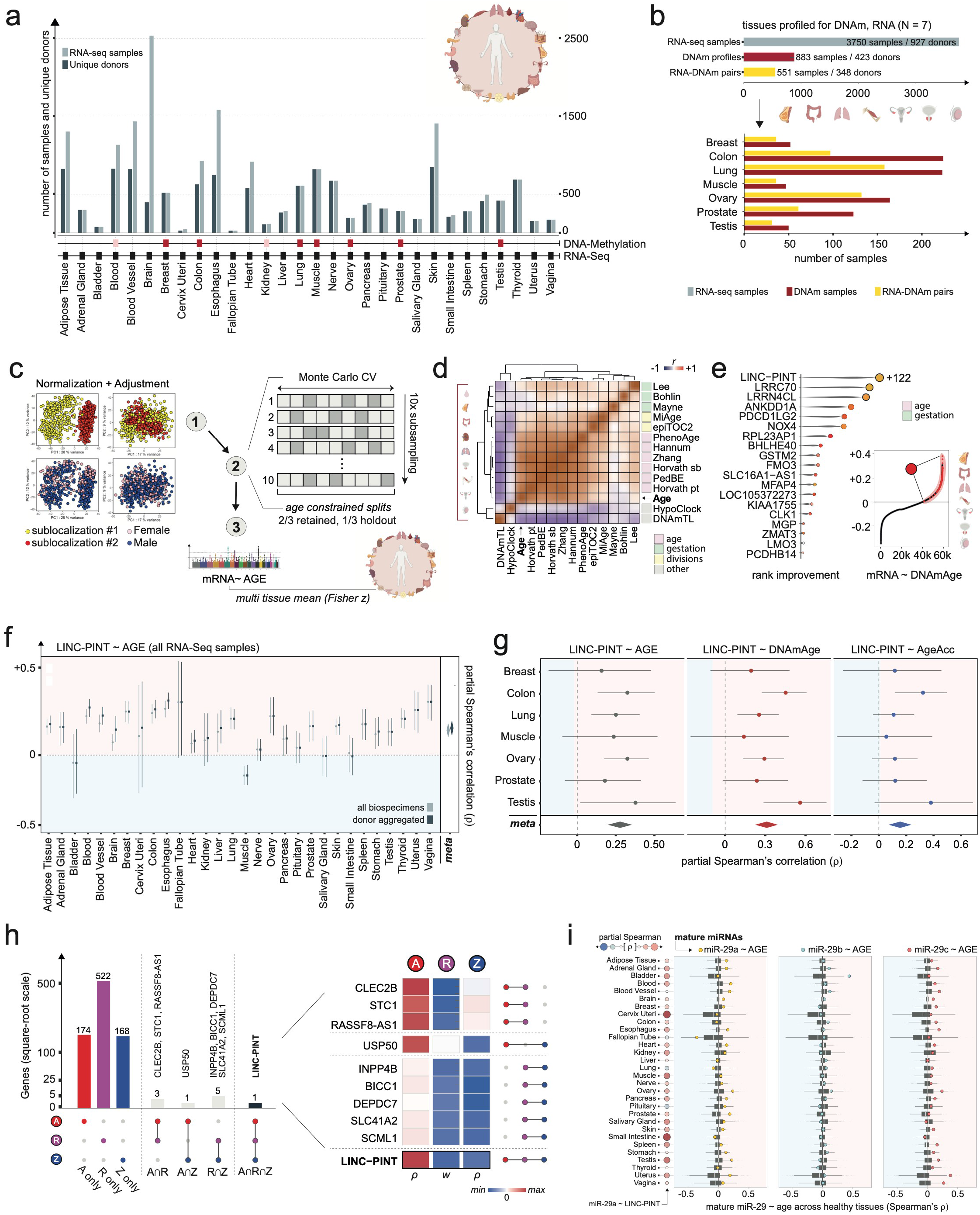
Coverage and sensitivity analyses supporting the discovery framework. **a**, RNA profiles and unique donors across 30 normal-tissue categories; tracks identify methylation availability and the seven paired-analysis tissues. **b**, Modality-specific availability in those seven tissues: 3,750 RNA records/927 donors, 883 methylation records/423 donors and 551 strict RNA–DNAm pairs/348 donors. Lower bars show per-tissue methylation profiles and strict same-tissue pairs; upper bars give modality totals. **c**, Normalization, covariate adjustment and age-constrained resampling workflow; illustrative projections are schematic. *Age-driven miR-29 trades youthful plasticity for epigenetic fidelity* **d**, Within-tissue clock/age correlation matrices averaged in Fisher-z space across nine tissues. **e**, Rank changes for the 177-gene seven-tissue chronological-age tail when DNAmAge replaces chronological age; *LINC-PINT* improves by 122 positions. The rank comparison uses the same tissue categories with endpoint-specific eligible samples, not strictly paired biospecimens. Grey spindles represent rank displacement, not distributions. **f**, Sex- and sublocation-adjusted *LINC-PINT*– chronological-age partial Spearman correlations, using all biospecimens or donor-aggregated expression. Whiskers are 95% donor-cluster bootstrap intervals (500 resamples); diamonds are descriptive random-effects summaries. **g**, *LINC-PINT* associations with chronological age, Horvath DNAmAge and acceleration in the 551 strict multimodal pairs. Intervals use 5,000 within-tissue donor bootstraps; diamonds summarize tissues. **h**, Exclusive intersections of the fixed A/R/Z signatures and the scores of genes in at least two sets. Heatmap columns have separate symmetric scales; opacity indicates membership, not significance. **i**, Covariate-adjusted *LINC-PINT*–miR-29a associations (first strip; full opacity, BH FDR < 0.05 across 30 tissues) and the positions of mature miR-29 family members within tissue-specific miRNA–age coefficient distributions. Boxes show medians and interquartile ranges; whiskers extend to 1.5 interquartile ranges. Some donors contribute to multiple tissues; pooled correlations are descriptive summaries. *Age-driven miR-29 trades youthful plasticity for epigenetic fidelity*

**Supplementary Fig. 2.**
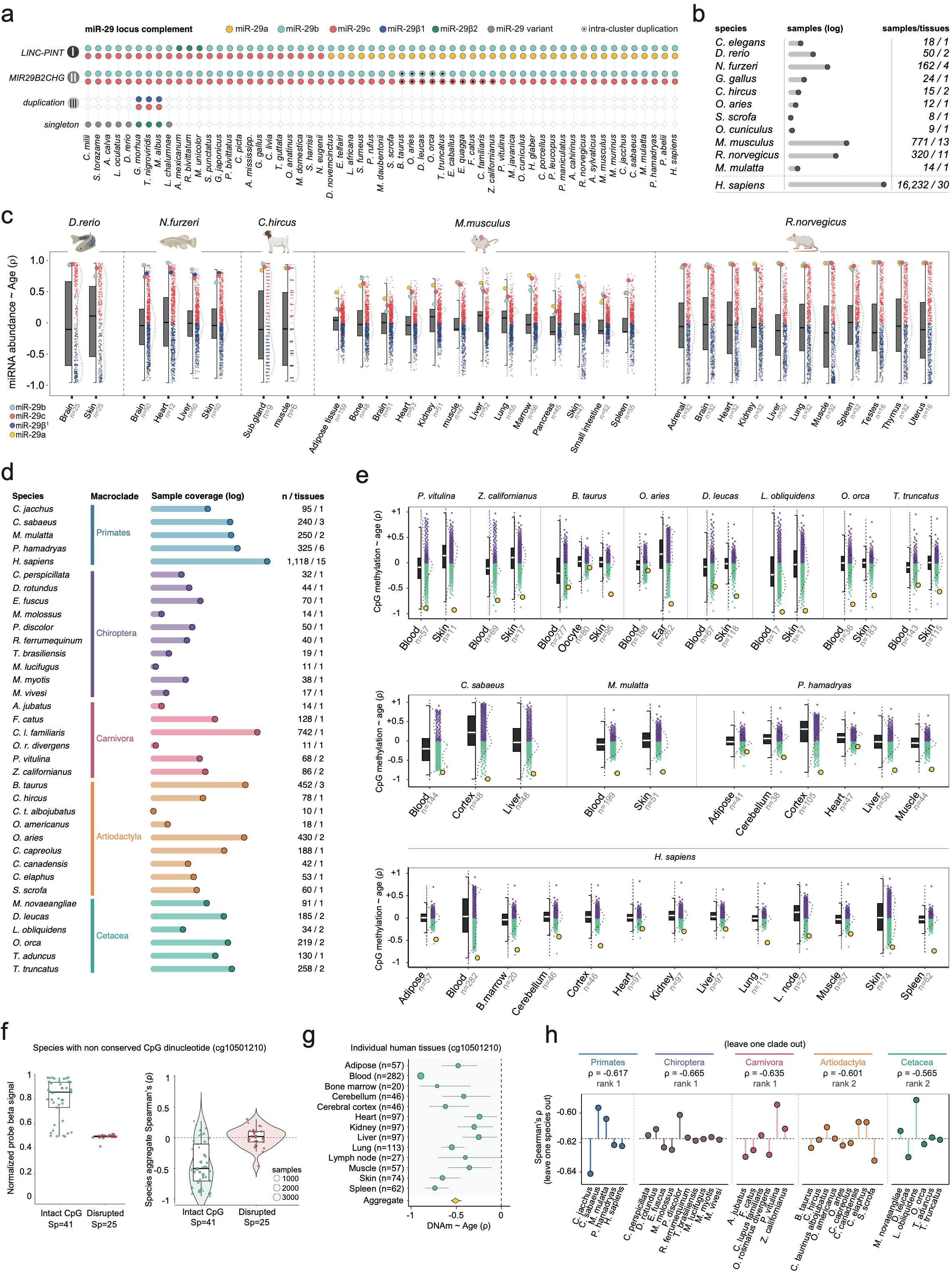
Tissue coverage and sequence-aware robustness of the cross-species analysis. **a**, Species-by-locus miR-29 complement. Colour denotes mature-sequence identity; marked copies indicate intracluster duplication. Empty positions denote missing annotations in the curated reconstruction. Sequence and locus reconstruction is described in Supplementary Methods. **b**, Expression-resource sample and tissue counts. The human row describes the Figure 1 GTEx RNA resource, not mature-miRNA libraries; humans do not enter the ten-species seed aggregate. **c**, Tissue-resolved miRNA–age Spearman distributions with miR-29 family members highlighted. **d**, Methylation coverage by species and analysis group after age-confidence and minimum-size filtering. **e**, cg10501210 within tissue-specific CpG –age correlation distributions; n denotes eligible profiles in each tissue. **f**, Species-level target-probe β signal and age association, contrasting conserved (41 species) and non-conserved (25 species) CpG targets. The latter include 19 disrupted dinucleotides and six genomes without an identifiable homologous probe-target region; 11 species without a suitable exact-species reference are excluded. These compare assay-target sequence contexts, not equivalent methylation measurements at an intact site. Point area represents sample coverage. **g**, cg10501210–age Spearman correlations in 13 human tissues with finite estimates. Whiskers are percentile 95% intervals from 2,000 within-tissue sample bootstraps. The diamond is the unweighted tissue-mean correlation with a corresponding bootstrap interval, not a random-effects meta-analysis. **h**, Leave-one-species-out and leave-one-group-out reaggregation of all 37,554 probes. Connecting segments indicate displacement from the full estimate, not confidence intervals. The target remains rank 1 after every species omission, including human, and rank 1 or 2 after group omission. Boxes in c, e and f show medians and interquartile ranges, with 1.5-interquartile-range whiskers; distributions describe variation among miRNA/probe coefficients or species.

**Supplementary Fig. 3.**
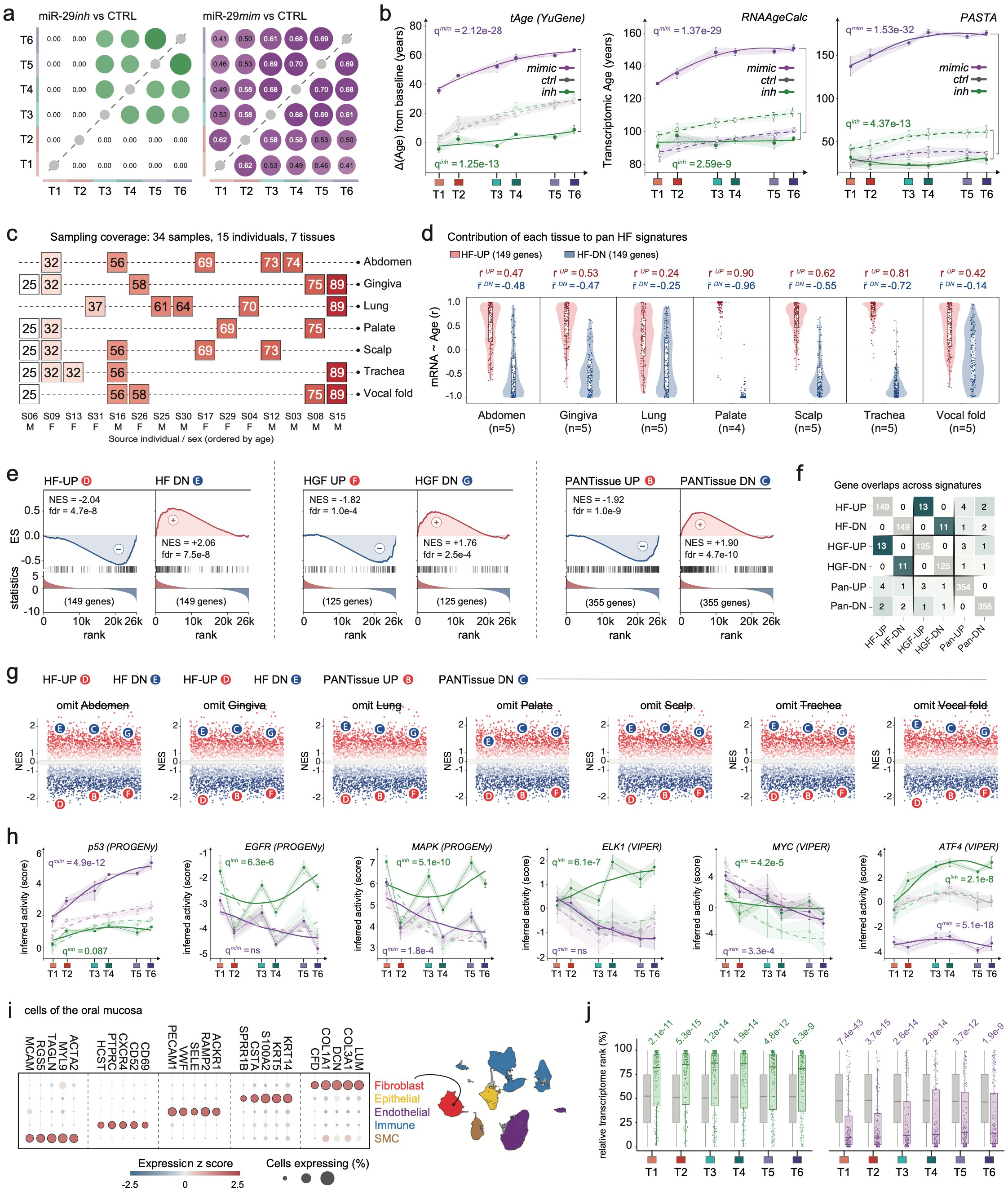
Control-resolved clocks and independent transcriptional ageing signatures. **a**, Jaccard overlap of differentially expressed gene sets across timepoints within each treatment arm. Up- and downregulated genes are combined; the statistic measures membership, not directional concordance. **b**, tAge, RNAAgeCalc and PASTA on their model scales, with both scramble-control arms resolved; q values test global shifts versus matched controls (four-test BH family per clock). **c**, External fibroblast cohort: 34 samples from 15 individuals and seven anatomical sites. **d**, Tissue-specific sex-adjusted Pearson age correlations of HF-UP and HF-DN genes; labels show arithmetic mean coefficients. Each tissue has five samples except palate (four). *Age-driven miR-29 trades youthful plasticity for epigenetic fidelity* **e**, Running enrichment for the six independently defined signatures B–G against the inhibitor-by-time ranking; labels report NES and multilevel BH FDR across all 4,721 tested sets, not only the six displayed signatures. **f**, Complete signature intersections (numbers) and Jaccard similarities (colour), not leading-edge overlaps. **g**, Pan-fibroblast signature enrichment after omitting each tissue; only D/E are rederived, while B/C/F/G remain fixed. **h**, PROGENy p53, EGFR and MAPK, and VIPER ELK1, MYC and ATF4 activities. **i**, Oral-reference marker expression and cell-class embedding; dot size is the fraction of expressing cells and colour the macrogroup-standardised mean expression. **j**, Signed-Wald transcriptome percentiles of miR-29 targets versus background genes. Higher percentiles denote more positive treatment-associated statistics. Significance uses a competitive CAMERA-style test, accounting for inter-gene correlation. Panels b and h show mean ± s.e.m. across five inhibitor/control and three mimic/control experiments; curves are descriptive fits. Global comparisons use matched-control, run-aware models. For h, BH correction covers 28 pathway–arm tests or ten transcription-factor–arm tests, including activities not displayed. Panel j boxes show medians/interquartile ranges and 1.5-interquartile-range whiskers; genes are not biological replicates.

**Supplementary Fig. 4.**
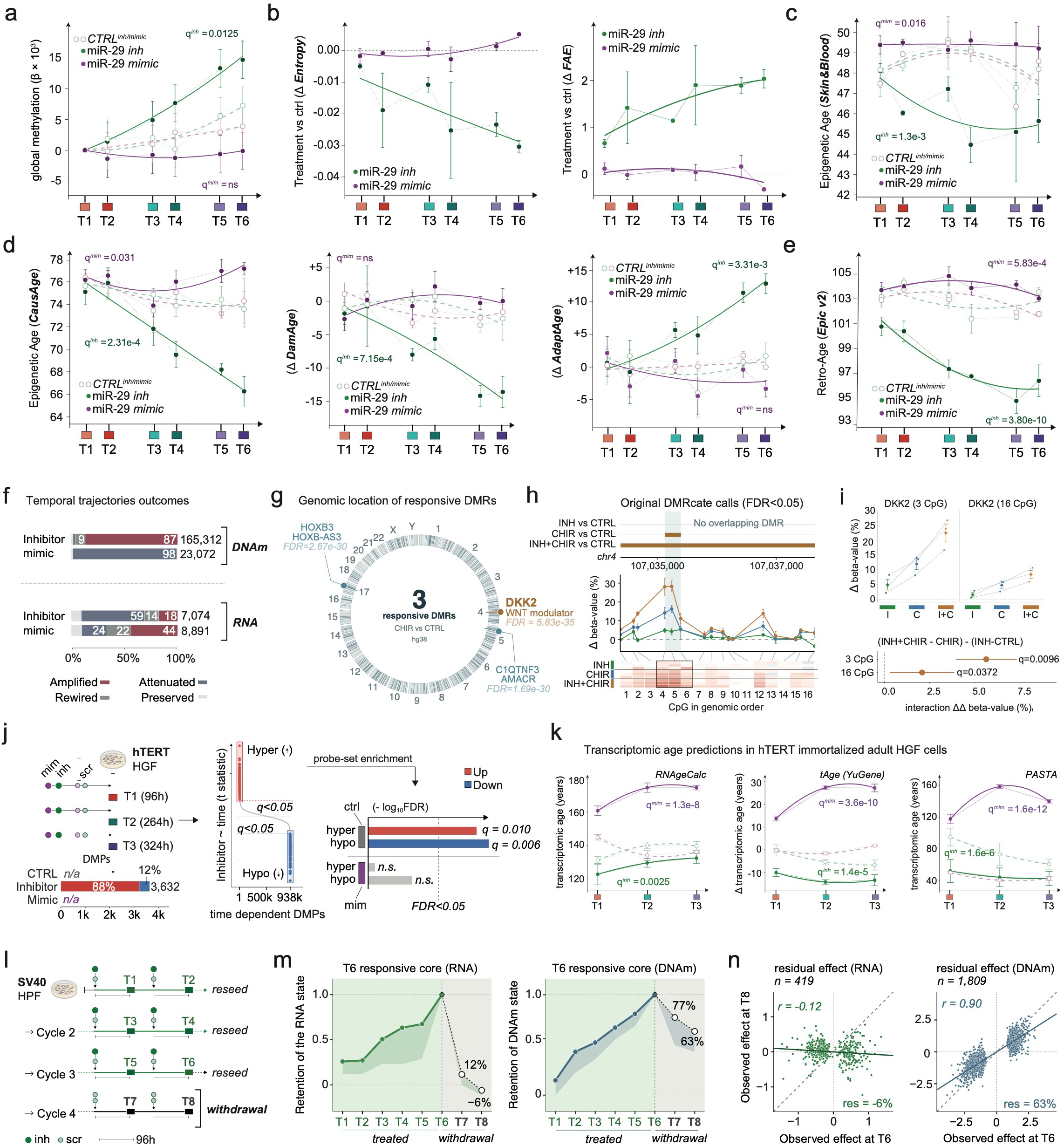
Epigenetic responsiveness to external stimulation and persistence after withdrawal. **a**, Global methylation with separate control arms; own-T1 changes and model-based 90% intervals; q values test treatment-by-time interactions versus matched controls. **b**, Entropy and FAE differences from same-experiment controls. **c–e**, Control-resolved Skin & Blood (c), CausAge, DamAge and AdaptAge (d), and RetroAge v2 (e). Points/bars show mean ± s.e.m. across three primary-HGF experiments per arm; global shift q values compare each treatment with its matched control (two-arm BH per clock). **f**, Temporal-feature classification for DNAm and RNA; categories combine slope direction, magnitude and nominal temporal support, not formal slope-difference tests. **g**, Three CHIR99021-responsive DMRs passing |mean Δβ| ≥ 0.10 among four native DMRcate calls. Labels report minimum smoothed FDR, not whole-region BH q values. **h**, *DKK2* DMR tracks, CpG-wise matched-control differences (mean ± s.e.m.) and experiment-resolved heatmap. Discovery includes incomplete EXP65; matched displays use complete EXP66–68. Connectors align genomic coordinates to CpG columns. **i**, Regional matched-control responses at the three-CpG CHIR99021 core and extended 16-CpG interval. Forest estimates are paired interactions, (INH+CHIR99021 − CHIR99021) − (INH − CTRL), on the beta scale, with 95% t intervals across three experiments; two-region BH q = 0.0096 and 0.0372. *Age-driven miR-29 trades youthful plasticity for epigenetic fidelity* **j**, hTERT-HGF design, temporal DMP counts, ranked statistics and competitive CAMERA enrichment of inhibitor-defined hyper/hypomethylating sets against control or mimic moderated t statistics. Bars show −log₁₀ BH FDR, not NES. **k**, hTERT-HGF transcriptomic clocks at three timepoints (four inhibitor/control and three mimic/control experiments; 42 RNA profiles; mean ± s.e.m.); q values test global shifts versus matched controls (four-test BH family per clock). **l**, SV40-HPF treatment/withdrawal design, with one inhibitor and one control sample at each of eight observations. **m**, Projection of matched effects onto the T6 response, normalised to one at T6. Shading spans seven nested RNA and five DNAm feature sets, not biological confidence intervals. **n**, Observed T8 versus T6 effects for 419 RNA and 1,809 DNAm features, with Pearson correlations and retained amplitudes. RNA and DNAm cores use nominal P < 0.05 and BH FDR < 0.05, respectively; the DNAm core-selection model includes post-withdrawal observations. HPF withdrawal is a single descriptive series; selected-locus *DKK2* inference is not independent of region discovery. The artwork label CHIR denotes CHIR99021.

**Supplementary Fig. 5.**
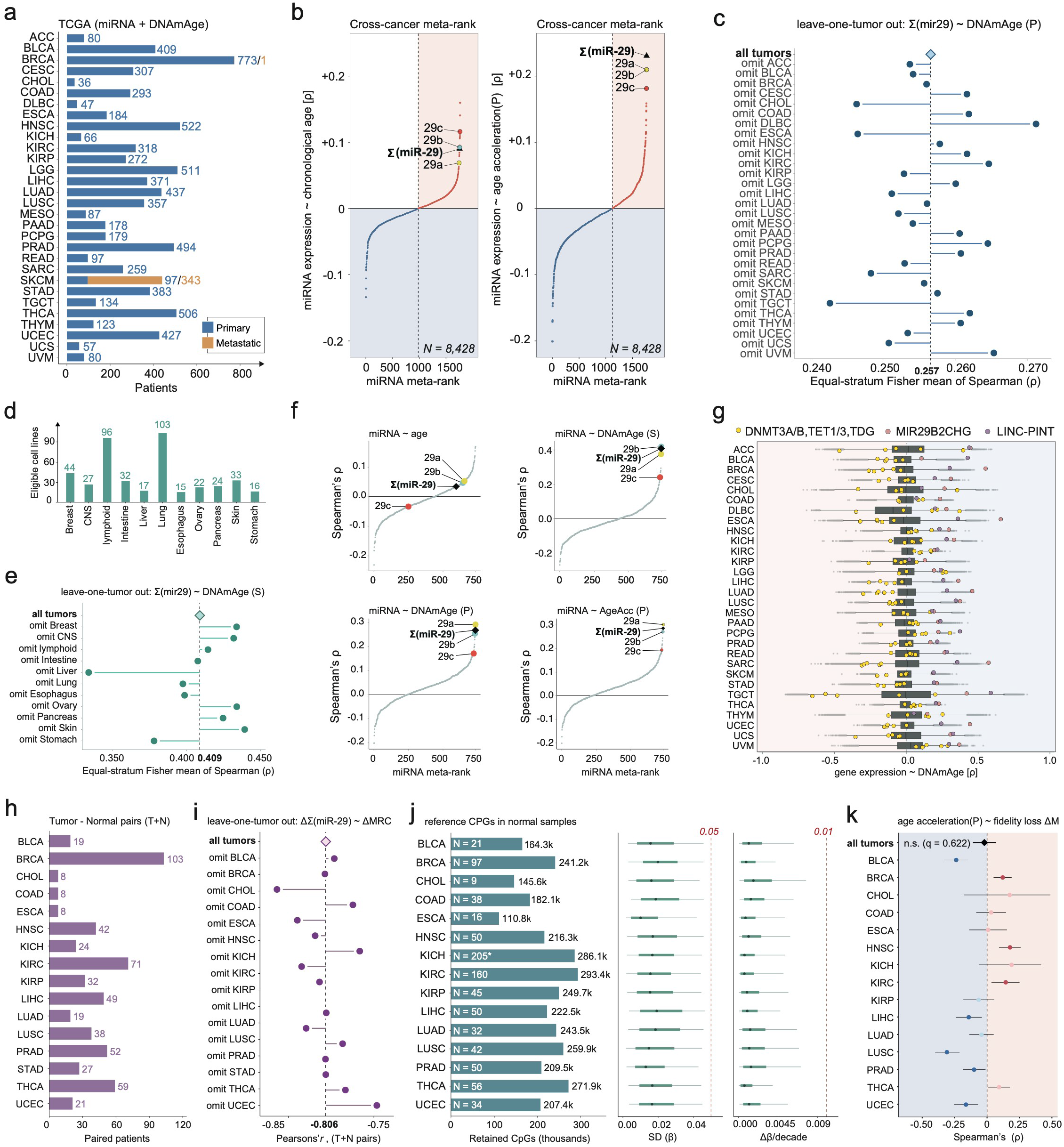
Cohort composition, age-readout sensitivity and normal-reference properties. **a**, TCGA miRNA–DNAmAge cohort: one tumour per patient, comprising 8,084 primary and 344 metastatic tumours. Metastases comprise one BRCA and 343 SKCM samples; adjacent normals are excluded. **b**, Equal-cancer miRNA rankings for chronological age and Horvath pan-tissue acceleration. **c**, Pan-miR-29–Horvath DNAmAge aggregate after leaving out each cancer. **d**, The 429 eligible CCLE lines across 11 lineages, each containing at least 15 complete cases. **e**, Leave-one-lineage-out pan-miR-29–Skin & Blood DNAmAge analysis. **f**, CCLE miRNA rankings for donor age, Skin & Blood DNAmAge, Horvath pan-tissue DNAmAge and Horvath pan-tissue acceleration. Ranked profiles in b and f are ordered coefficients, not fitted response curves. **g**, Cancer-specific RNA–Horvath DNAmAge coefficient distributions, highlighting the five MRC genes, *LINC-PINT* and *MIR29B2CHG*. Boxes show medians and interquartile ranges with 1.5-interquartile-range whiskers. **h**, Counts of strict paired tumour–normal patients with both RNA and miRNA assays (580 patients, 16 histotypes). **i**, Pearson correlation between cancer-mean paired miR-29 and MRC effects after each cancer omission. Connectors in c, e and i show changes from the full estimate, not confidence intervals. **j**, Retained CpGs and normal-reference sample counts; KICH uses 205 shared kidney normals. *Age-driven miR-29 trades youthful plasticity for epigenetic fidelity* Stability summaries show medians, interquartile ranges (thick) and 5th–95th percentiles (thin); dashed thresholds are SD(β) = 0.05 and |Δβ| per decade = 0.01. **k**, Age-spline-adjusted association between Horvath acceleration and reference deviation: approximate 95% correlation intervals and REML/Knapp–Hartung summary (pooled ρ = −0.0208, 95% interval −0.1088 to 0.0676; q = 0.622). Absence of a pooled association does not imply equivalence of these readouts within every cancer.

