## supplementary methods for "Age-driven miR-29 trades youthful plasticity for epigenetic fidelity"

Statistical procedures and selected estimates are described below.

### 1. Public-data discovery, matching and aggregation

Reprogramming analyses used unstranded STAR gene counts from GSE165176 after removing Ensembl version suffixes. Samples were restricted to experiment 1 and days 11, 13 and 15. The six donors, N2, N3, O1, O2, Y1 and Y2, were aged 0, 0, 53, 53, 29 and 35 years, respectively. The separate methylation trajectory comprised 22 profiles from the four adult donors at days 9, 11 and 15. The three-standard-deviation negative-tail cutoff was calculated over finite DESeq2 Wald statistics.

Embryonic RNA cohorts were combined at the count level. Variance stabilization included developmental group and GEO cohort; limma batch correction preserved developmental-group effects. Slingshot inferred a single lineage, oriented toward increasing developmental stage. The 175-gene lower tail was defined from Fisher-transformed gene–pseudotime Spearman coefficients, independently of the P values displayed on the rank plot. Heatmap rows were standardised, with the colour scale limited to ±4; samples were ordered by pseudotime.

The GTEx RNA atlas covered thirty normal tissue categories. Strict same-biospecimen matching yielded 551 RNA–methylation pairs from 348 donors across seven tissues. Exact donor ages came from controlled-access annotation.

The ageing signature used precomputed tissue-level association coefficients from analyses with normalization, sex and anatomical-sublocation adjustment, and repeated age-constrained subsampling retaining two-thirds of samples. The cross-tissue tail threshold was mean(z) + 3 SD(z), where z is the aggregated Fisher-transformed association. The three gene sets were intersected by Ensembl identifier.

A complementary cutoff-free analysis combined directional ranks across 34,686 jointly assessed genes from the same datasets. R and Z were ranked from lowest to highest and A from highest to lowest. Ranks were divided by the number of genes, with average ranks for ties; their geometric mean defined the score, with smaller values indicating stronger concordance. *LINC-PINT* ranked first.

*LINC-PINT* tissue-specific analyses used partial Spearman correlations: expression and the age-related predictor were ranked and residualised on estimable sex and anatomical-sublocation covariates. Repeated biospecimens were either retained with donor-cluster resampling or averaged within donor and tissue before correlation. The thirty-tissue percentile intervals used 500 donor-cluster bootstrap draws. Strictly paired RNA–DNAm analyses used 5,000 donor bootstrap draws, with age acceleration defined as the within-tissue OLS residual of Horvath DNAmAge on chronological age. The acceleration association additionally adjusted for ranked age. Cross-tissue

REML summaries used bootstrap variances on the Fisher-z scale; these summaries did not account for all covariance from donors contributing multiple tissues.

Healthy-tissue mature-miRNA coefficients came from the processed GTEx v10 small-RNA resource (17 March 2025 release; dbGaP phs000424; https://gtexportal.org/home/), incorporating the consortium's miRXplore-contamination correction. RNAcentral v19 identifiers and annotations were retained.

The healthy-tissue mature-miRNA summary used the arithmetic mean of tissue coefficients. Published developmental expression data from Russell et al. [55] were log1p-transformed and related by Pearson correlation to the stage sequence (1,2,3,4,5,6,4,5,6). Reprogramming miRNA data came from GSE62776. *MIR29B* locus abundances were summed, counts were converted to log counts per million, and Pearson correlations used days 0,2,5,8,10,14,20,24 and 25.

### 2. Comparative sequence and age-association analyses

The vertebrate tree was retrieved from Open Tree using rotl [56]; unresolved branching points were converted to bifurcations using ape with a fixed, non-random rule [57]. Cluster architecture, host-locus annotations and sequence histories were curated from accession-level records. Missing annotations did not establish biological absence. Comparative miR-29 copy labels identify sequence groups rather than human *MIR29B1* and *MIR29B2* genomic copies. Motif annotations describe sequence features, not measured localization in each species.

The vertebrate catalogue comprises 59 species, 121 clusters and nine singleton loci, with 257 clustered precursor fragments and nine singleton fragments (266 loci in total). Genomic positions were assigned by exact matching to the specified reference chromosome or scaffold on either strand, with the mature sequence located within its precursor fragment. Coordinates are one-based and inclusive; sequences are reported 5′ to 3′ in RNA orientation. Precursor entries are genomic fragments surrounding the mature sequence, not experimentally mapped processing boundaries. Intracluster copies are ordered from the first miR-29b-like locus along the strand-aware cluster orientation; the distal repeated member is designated the duplicate.

Raw small-RNA reads were reprocessed for Caenorhabditis elegans (GSE110846, GSE110849), Danio rerio (GSE117807, GSE117820), Nothobranchius furzeri (GSE104321 [44], GSE107062 [36]), Gallus gallus (GSE139304 [58]), *Capra hircus* (GSE166842 [59], PRJNA553597 [60]), Ovis aries (PRJNA1195695 [61]), Sus scrofa (GSE64275 [62], PRJNA196673 [63]) and Oryctolagus cuniculus (PRJNA692651 [64]). The *C. elegans* and D. rerio cohorts were obtained from the JenAge GEO deposits. The workflow used nf-core/smrnaseq v2.4.1 with Nextflow v25.04.8 [65] and Singularity v3.8.0 [66]. Species-specific mature and hairpin FASTAs came from the May 2026 miRBase reference snapshot. Reference genomes were WBcel235 (*C. elegans*), danRer11 (D. rerio), Gallus_gallus-5.0 (G. gallus), CHIR_1.0/GCF_000317765.1 (C. hircus), Oar_v4.0 (O. aries), Sscrofa11.1/GCF_000003025.6 (S. scrofa) and OryCun2.0/GCF_000003625.3 (O. cuniculus).

Adapter trimming and read-length processing used fastp. Chicken used adapter autodetection with minimum and maximum read lengths of 18 and 30; the other reprocessed species used the 3′ adapter TGGAATTCTCGGGTGCCAAGG with settings 20 and 35. The maximum-length setting capped read length rather than excluding longer reads. Contamination filtering and miRDeep discovery were disabled. Mature-miRNA counts came from Bowtie/SAMtools quantification.

For N. furzeri, mature and hairpin sequences were extracted from the NFINgb Nfu_20150522 soft-masked genome and matching miRNA GFF3 annotation (https://nfingb.leibniz-fli.de/), using one-based inclusive coordinates, reverse complementation on the negative strand and T-to-U conversion. Predicted identifiers and source arm labels were retained for sequence-based seed assignment.

Mouse (GSE217458 [28]), rat (GSE172269 [67]) and macaque (GSE18013 [68]) analyses used published expression matrices: normalised reads per million for mouse and expression/count tables for rat and macaque. Source-specific transformations and tissue aggregation were retained. Public dataset accessions, source publications and reprocessing settings are described in this section.

The age of C. hircus sample SRR9663090 was set to one day. The *Macaca mulatta* cohort was restricted to fourteen brain profiles, excluding testis. The seed analysis combined mature miRNAs sharing a seed within species in Fisher-z space, then species within clades, and finally five equally weighted groups. It retained 76 seeds represented in at least one species of each group and excluded human expression profiles.

The miR-29 seed AGCACCA ranked first among the 76 eligible seeds (aggregate $\rho = 0.8923$). Its rank remained first after every single-species omission ($\rho = 0.874$–$0.906$), single-group omission ($\rho = 0.874$–$0.904$), equal-species aggregation ($\rho = 0.870$) and $(n-3)$-weighted aggregation ($\rho = 0.743$).

Mammalian-array analysis retained 37,554 probes. Age-confidence filtering preceded within-species tissue eligibility. Strata with invariant age annotations, including human keratinocytes and placenta, had no estimable age correlation and were excluded. Tissue coefficients were averaged arithmetically within species; Fisher averaging was used above species. *Myotis* and *Tursiops* were represented as balanced genus-level units where specified by the curated hierarchy. Species omissions rebuilt the corresponding genus unit, all higher-level means and the full probe ranking. Clade omissions equally weighted the four remaining clades.

The cg10501210 conservation inventory contains 77 species with at least one species–tissue stratum containing ten or more samples after excluding missing ages and age-confidence scores ≤80. Eleven species lacked a suitable exact-species genome and were excluded from conservation comparisons. Of the remaining 66, 60 had an identifiable homologous target region: 41 retained the CpG and 19 had a disrupted dinucleotide. Six species with an available genome but no identifiable homologous region were also classified as non-conserved, giving 25 non-conserved targets. The alignment therefore contained the 60 species with available sequences. The five-group methylation ranking used 36 conserved-target species in the specified clades. Cross-species sequence checks assessed retention of the CpG dinucleotide corresponding to human cg10501210; comparator probes were not individually screened.

For Fig. 2i, the 60 target-region sequences were submitted to the PhyML-SMS/OneClick workflow at NGPhylogeny.fr [46]. MAFFT v7.467, using L-INS-i, generated the multiple alignment [69], which was trimmed with BMGE [70]. The BMGE-cleaned FASTA was displayed in MSAViewer with the nucleotide colour scheme [71]. The displayed panel illustrates the target-region alignment. The downstream phylogenetic-tree output was not used to define CpG conservation.

For the human cg10501210 forest, 2,000 within-tissue bootstrap draws resampled paired age and beta observations; intervals used the 2.5th and 97.5th percentiles. The overall estimate was the unweighted arithmetic mean across thirteen estimable tissues, with an interval from the mean tissue coefficient in each draw. This interval is conditional on the included tissues and does not account for shared donors across tissues. The sequence-status comparison used 41 conserved and 25 non-conserved targets, averaging normalised beta first within tissue and then across tissues. Non-conserved targets served as assay-context controls and were excluded from the conserved-site ranking; their signals do not measure methylation at an intact homologous CpG. This descriptive comparison did not adjust for phylogenetic relatedness.

#### 3. Growth, transcriptional signatures and regulatory activity

Growth measurements were normalised within replicate as 100 × (count−count$_{T0}$)/(maximum count−count$_{T0}$). Means and SEM were calculated across three experiments per arm, and curves were exponential-plateau fits to replicate measurements. This normalization describes relative growth trajectories, not population-doubling times.

RNA PCA used variance-stabilized expression after sequential adjustment for experimental arm and batch while preserving the experimental design. Gene centering and scaling preceded PCA. Global DESeq2 treatment-versus-own-control tests used BH adjustment across 20,982 inhibitor-arm and 24,198 mimic-arm genes passing independent filtering. Within each arm, Jaccard overlap used genes with finite adjusted P values and fold changes at all six timepoints. For each timepoint pair, overlap was the number of genes significant at both times divided by the number significant at either time, regardless of effect direction. Comparisons between two empty sets were undefined.

External fibroblast age correlations used VST expression from GSE140523 [47]; age and expression were separately residualised on sex with an intercept. Correlations were clipped to $\pm(1-10^{-6})$ before Fisher transformation. Pan-fibroblast aggregation required at least five finite tissue correlations, including in leave-one-tissue-out analyses. Tail thresholds were computed in the entire estimable universe before excluding the two miR-29-associated host transcripts. The gingival universe contained 12,467 genes, giving 125 in each strict tail. Each tissue had five samples except palate, which had four.

Enrichment used 25,987 signed Wald entries for the inhibitor-by-time term. The ranking retained 57 duplicated gene-symbol entries; gene-set matching used the first occurrence. Plotted NES were calculated with fgseaSimple version 1.24.0 using 10,000 permutations, minimum matched size 20, no upper size bound, gseaParam = 1 and scoreType = std. Significance used fgseaMultilevel with sampleSize = 501, nPermSimple = 10,000 and eps = 0. BH correction covered all 4,721 tests separately in each analysis, including omission analyses.

PROGENy used the human model, the top 500 genes per pathway, 10,000 permutations and z-score output on within-arm, batch-adjusted VST expression. VIPER used the DoRothEA regulon through decoupleR to obtain per-sample scores [72]. Treatment-minus-own-control means were calculated at each timepoint. Overall-shift models included categorical time and a run random intercept; BH correction covered ten transcription-factor–arm tests or 28 pathway–arm tests, including undisplayed activities. Heatmap clustering used Euclidean distances and complete linkage without prior row z-scoring; columns retained time order. Competitive miR-29 target enrichment used CAMERA-style testing with assumed inter-gene correlation 0.01.

The oral single-cell reference contained 133,335 cells. The fibroblast macrogroup combined homeostatic and inflammatory/remodelling subsets. Bulk projections included all 96 experimental RNA-seq samples. Marker dotplots used cell-count-weighted normalised expression; colour represented the across-macrogroup z-score of log1p mean expression, and point area represented detection frequency. The marker summary included five macroclasses containing 131,475 cells; cycling/myogenic categories remained only in the embedding.

**4. Primary-HGF molecular-age predictions and inference**

Primary-HGF RNAAgeCalc predictions retained their native age scale. tAge used a human-inclusive YuGene model, with per-gene median control expression across time defining an arm-specific input reference. The main display used own-arm T1 control recentering, a 122.5 multiplier and postprediction batch adjustment preserving treatment-by-time groups. Supplementary curves retained unadjusted predictions with controls separated. PASTA applied its model-gene filtering and rank normalization once before prediction.

Skin & Blood predictions used meffonym 1.2.1 with fixed calibration parameters and adjusted, collapsed beta values. RetroAge used its intercept and 1,378 available EPIC v2 assay coefficients on adjusted noncollapsed beta; missing entries were replaced by the corresponding probe's cohort mean. CausAge, DamAge and AdaptAge had 496/585, 1,016/1,089 and 938/999 available features, respectively. Absent features were omitted without renormalization and represented 13.8%, 6.8% and 1.6% of absolute coefficient weight. The resulting scores were interpreted as relative model responses. Skin & Blood and Horvath pan-tissue coverage was 374/391 and 340/353 features. Six DamAge entries and one AdaptAge entry were imputed using the probe's cohort mean. DamAge and AdaptAge plots used a constant pooled-control-T1 offset while preserving score direction.

CausAge, DamAge and AdaptAge were originally trained to predict chronological age, but their biological interpretations differ. Higher AdaptAge acceleration was associated with lower mortality, and AdaptAge increased during reprogramming in the original study [18]. An increase in AdaptAge therefore does not by itself imply an adverse response or contradict a decrease in DamAge.

Clock plots show group mean ± SEM unless specified otherwise; ribbons summarize pointwise uncertainty, and quadratic curves are descriptive fits. Ordinal display coordinates 1,2,4,5,7,8 group paired timepoints and do not represent elapsed hours.

Primary-HGF RNA-clock global shifts used maximum-likelihood comparisons of score ~ factor(timepoint) + (1|run) with score ~ treatment + factor(timepoint) + (1|run). Fig. 3e compares each treatment with all eight control runs, whereas Supplementary Fig. 3b retains own-arm controls. Inference used native predictions with unchanged arm-specific tAge input normalization. BH adjustment was performed separately for each control definition across four tests per clock: two arm-specific shifts and two condition-by-quadratic-time interactions. Timepoint tests used Kenward–Roger degrees of freedom and Holm correction across six contrasts per arm and clock.

Primary-HGF epigenetic-clock global comparisons used Kenward–Roger F tests for average treatment shift, controlling categorical time and a run random intercept. BH correction covered the two arms separately for each control definition. Main curves use pooled controls and supplementary curves matched controls; both display global q values only. The twelve timepoint contrasts per epigenetic clock formed a separate testing family.

#### 5. Methylation distributions, compartments and temporal geometry

Preprocessing retained both assay-suffixed EPIC v2 identifiers and base-CpG identifiers. For PCA, noncollapsed M values were corrected sequentially for experimental arm and batch, preserving the time-by-group design. Of the 20,000 most variable features, 19,876 with finite measurements entered the standardised PCA. The global-mean model used raw collapsed beta values from CpGs measured in every sample.

Global methylation was fitted using condition-by-categorical-time means and an experimental-run random intercept. Contrasts subtracted each condition's own T1 mean. The two control arms were equally combined in the main display and pooled-control comparisons; supplementary curves retained matched comparisons. Intervals were model-based 90% confidence intervals. Treatment-by-time interaction was tested while allowing a constant treatment offset, with BH correction across the two arms. Entropy and fraction-at-extremes T6 comparisons used mixed-model t contrasts with Kenward–Roger degrees of freedom. BH correction covered four metric-by-arm tests separately for pooled-control and matched-control comparisons.

Beta densities used adjusted noncollapsed values bounded to [0,1], a kernel bandwidth of 0.02 and 512 evaluation positions without boundary reflection. Curves summarised mean sample density and, where shown, between-sample SD. Entropy was $-\Sigma\, p_k \log(p_k)$ over twenty equal-width beta bins;

FAE was the fraction of beta values below 0.2 or above 0.8. GTEx entropy and FAE were tissue-adjusted, summarised in fifteen-sample windows advanced by five samples, and retained in age bins supported by at least six windows. The 161 overlapping windows support descriptive smoothing, not independent window-level tests. Upper panels used DNAmAge; the lower beta-density comparison used chronological age.

CpG context was assigned with island > shore > shelf > open-sea priority. Promoter annotations combined TSS1500, TSS200, 5′ UTR and first exon; promoter-CGI and island categories overlap. Net temporal burden was 100 × (significant hyper−significant hypo)/tested probes, and point size encoded 100 × (hyper+hypo)/tested probes. Repeat annotation used the hg38 RepeatMasker resource AH99003 with strand-aware overlaps. The ERVL category combined ERVL and ERVL-MaLR; DNA and SINE summaries covered the whole classes, including hAT and Alu, respectively. Compartment and repeat heatmaps were row-standardised over the eighteen group–time combinations.

Control-selected temporal movers comprised 2,188 hypomethylating and 20,884 hypermethylating assay probes. Values were centred on each condition's T1 mean, averaged within run and fitted with GAMs containing a four-dimensional time smooth and a random-run term. Prediction excluded the run-specific smooth and used 90% intervals. Categorical-time interaction inference was separate from those descriptive fits, with four direction-by-arm tests in its BH family. Descriptive trajectory classes required nominal support for an individual slope; loss of support was distinguished from a demonstrably smaller slope. Supported same-sign slopes were compared using a 20% magnitude band.

Slope comparisons used through-origin regression on separate unions of treatment- and control-significant probes. The inhibitor union contained 165,312 probes, of which 97.4% were inhibitor-significant; the mimic union contained 23,072 control-significant probes because no probe met the mimic threshold. The coefficients 2.389 and 0.0707 describe different selected sets and are not directly comparable amplification estimates. The separate control-selected trajectory analysis applied the same control-defined probe set to both treatments; the control observations were used both to select that set and to estimate its response.

For the methylation-displacement representation, condition/time centroids used the 50,000 most variable finite adjusted M-value features, selected by total variance across all samples. Consecutive centroid differences defined step vectors. Their Gram matrix, divided by the number of features, supplied a common two-dimensional eigenspace. Each projected vector was rescaled to its original RMS magnitude and accumulated from a common origin, preserving step lengths but not every high-dimensional distance.

#### 6. Fibroblast-reference deviation and stimulus-responsive methylation

The 123-CpG reference was constructed in three stages using external data. Of 485,512 measured GSE77135 probes, 384,494 autosomal, platform-compatible CpGs entered the stability screen. Age and sex assessments used donor means. The cohort comprised ten male donors and one female donor, so the sex-effect bound was a screening constraint rather than strong evidence of sex independence. The reference centroid was the median beta at each retained CpG. GSE179847 filtered predicted culture-associated change before and after batch adjustment. GSE31848 repeated arrays were averaged by cell line before a limma contrast with equal weights for epithelial and smooth-muscle/perivascular groups and robust empirical Bayes. Selection required sex-adjusted BH q ≤0.20, |delta-beta| ≥0.10 in adjusted, unadjusted and purified-immune contrasts, concordant direction and the stability criteria specified in Methods. These thresholds defined operational stability; statistical equivalence was not tested. Both methylation directions were eligible. The immune cohort contributed to selection rather than held-out validation.

HGF reference-deviation scores used beta values without batch correction and were compared with same-run, same-time, own-arm controls. Cycle centers were 84, 180 and 276 h. The central display summarizes three within-run cycle means with a descriptive quadratic and interpolated SEM; its q value tests mean matched displacement over T1–T6. One beta percentage point corresponds to 0.01 in average absolute beta distance. The 123-CpG reference was annotated for hg38 position, direction, CpG context, genes and assay SNPs; these SNP annotations do not establish sample genotypes.

The inhibitor-minus-control mean difference over T1–T6 was 0.379 beta percentage points (three run means; 95% t interval 0.194–0.564; two-sided P = 0.01263, two-arm BH q = 0.02526). The late T5/T6 difference was 0.581 percentage points (95% interval 0.377–0.784; P = 0.00658, q = 0.01316). Mimic effects were −0.012 and −0.105 percentage points, respectively (P = q = 0.89538 and 0.42385). Each BH family covered the two arms within one temporal summary. Duration-weighted summaries gave inhibitor effects of 0.361 percentage points for the cycle-mean curve (P = 0.01236, q = 0.02473) and 0.391 for the full trajectory (P = 0.01737, q = 0.03474); these assess average displacement, not temporal slope.

Additional post hoc reference-deviation checks were performed. A one-sided t test on the same three late run means gave inhibitor P = 0.00329 and two-arm BH q = 0.00658. This analysis was not prespecified and did not replace the two-sided test; its correction did not account for selecting among methods. Exact signed-rank and sign-flip tests gave inhibitor P = 0.250 two-sided and P = 0.125 one-sided, reflecting their limited resolution with three runs. A Welch comparison between matched-control treatment effects gave an INH-minus-MIM difference of 0.686 percentage points (95% interval 0.302–1.069; P = 0.0118). Timepoints and CpGs were not treated as independent biological replicates.

The global-shift diagnostic calculated each matched pair's mean raw beta difference over 924,875 collapsed CpGs measured in every sample. This scalar was added to each control reference-site beta before recalculating distance from the unchanged centroid; a complementary analysis subtracted it from treated beta values. At T5/T6, imposing the inhibitor global shift of +0.962 beta percentage points uniformly predicted a distance change of −0.050, compared with the observed +0.581; recentering retained +0.723. The observed-minus-counterfactual difference was +0.631 (95% interval 0.278–0.985; P = 0.0165), and the recentered effect had a 95% interval of 0.450–0.996 (P = 0.00760). Two-arm BH q values were 0.0331 and 0.0152; adjustment across all 76 diagnostic tests gave 0.0993 and 0.0892. A uniform mean shift therefore did not reproduce the observed displacement; this does not exclude heterogeneous remodelling or establish functional identity loss.

Direction-resolved summaries used the externally defined 36 fibroblast-high and 87 fibroblast-low sites. Late inhibitor distance increased by 1.236 and 0.309 beta percentage points, contributing 0.362 and 0.219 to the all-site score; raw beta increased in both subsets. Multiplying each matched beta difference by the sign of the external cell-type contrast gave a signed fibroblast-oriented margin change of −0.031 percentage points (95% interval −0.841 to 0.779; P = 0.884). Increased absolute distance therefore did not establish a coherent shift toward another lineage.

Genomic annotation places one selected CpG near *CCN1*, four near *DNM3OS* and one near *FAP*, representing three loci. These annotations provide biological context, without establishing gene regulation or functional enrichment.

CHIR99021 DMRcate analysis used ~0 + GROUP + BATCH on M values, with BATCH identifying biological experiment, IlluminaHumanMethylationEPICv2 annotation 20a1.hg38, epicv2Remap = FALSE, CpG FDR = 0.10, lambda = 1,000 and C = 2. Untreated ZERO samples entered discovery; factorial regional comparisons used matched controls from the three complete experiments, EXP66–68. Regions were called separately for each comparison. min_smoothed_fdr is the minimum smoothed evidence within a region, not a whole-region BH q value. Three CHIR99021 loci passed |mean delta-beta| ≥0.10. *DKK2* was selected for WNT-pathway relevance, with a three-CpG core nested within a sixteen-CpG interval. The paired contrast was 100 × [(INH+CHIR99021−CHIR99021)−(INH−CTRL)] on regional mean beta, with 95% t intervals across three experiments and two degrees of freedom. Interactions were 5.63 and 2.09 beta percentage points for the core and extended intervals (two-region BH q = 0.0096 and 0.0372). CpGs were averaged before inference. Correction covered the two nested intervals, not selection of *DKK2*; discovery and inference shared experiments, so the selected-locus result was not independently validated.

### 7. Additional fibroblast models

#### hTERT-immortalized gingival fibroblasts

The hTERT-HGF model (ABM T0026; hTERT-hNOF, RRID:CVCL_C0UE) was established from primary gingival fibroblasts of a healthy adult [38]. Supplier documentation identifies the line as male-derived; exact donor age was not reported. RNA profiling comprised four inhibitor/control and three mimic/control runs at three timepoints (42 samples).

RNAAgeCalc used FPKM adjusted sequentially for experiment and batch while preserving group and time effects; negative adjusted expression was set to zero before applying the all-tissue Caucasian model. tAge used version 1.1.0 preprocessing and its predictor under scikit-learn 1.3.2, with arm-specific control subtraction. Its display used own-arm T1 control recentering, the 122.5 multiplier and batch adjustment preserving arm means. These outputs describe relative model responses rather than elapsed culture age. Matched-control global inference used likelihood-ratio tests fitted by maximum likelihood, with BH correction across two shifts and two interactions per clock. Six timepoint contrasts per clock formed a separate testing family.

Temporal methylation analysis tested 937,687 assay probes. Inhibitor-derived hyper/hypomethylating sets were evaluated against control and mimic temporal moderated t statistics using limma cameraPR with use.ranks = FALSE and inter.gene.cor = 0.01. BH correction covered two sets within each target condition. Bar heights were $-\log_{10}$ FDR and colours indicated CAMERA direction.

Reference-deviation analysis applied the same 123 sites and external centroid to 36 methylomes: three treatment/control runs per arm, sampled 96 h after each of three treatment cycles. EPIC v2 replicate assays were collapsed on the beta scale; all 123 CpGs had complete measurements. Treatments were compared with their contemporaneous own-arm controls within run. The final comparison used the third-cycle 96-h harvest. The inhibitor effect was +0.122 beta percentage points (95% t interval 0.046–0.198; P = 0.02046, two-arm BH q = 0.04093), with all three runs positive; mimic P = q = 0.40590. Averaging all cycles within run gave an inhibitor effect of +0.133 percentage points (95% interval 0.017–0.249; P = 0.03885, q = 0.07769), with mimic P = q = 0.29397. Two-sided one-sample t tests used three run-level matched differences per arm, with separate two-arm BH families for final-cycle and full-series comparisons.

#### SV40-immortalized pulmonary fibroblasts

The SV40-HPF model (ABM T0490) was derived from a 65-year-old female donor. Its DNAm series comprised sixteen samples, one inhibitor and one control at each of eight observations. T1–T6 were 96-h post-treatment collections; T7 and T8 were 168 and 240 h after withdrawal. Probe selection used M values collapsed after M conversion and a piecewise model of matched differences: T6 offset, active slope (−5 to 0 at T1–T6, then zero) and post-withdrawal slope (0 during treatment, then 168/96 and 240/96). limma empirical-Bayes tests used trend = FALSE and robust = FALSE, with BH correction within contrast. The 1,809-site core was selected by T6-offset FDR <0.05 using all observations, including those after withdrawal. Projection used M values calculated from collapsed beta with this fixed core. RNA used a 419-gene core selected by nominal T6 P <0.05 and observed VST differences.

For each fixed feature universe, retained amplitude was $R_t = \Sigma_j \delta_{j,T6} \delta_{j,t} / \Sigma_j \delta_{j,T6}^2$, with T6 equal to one by construction. Envelopes span seven nested RNA universes and five DNAm universes, giving T8 amplitudes of −11.4% to −6.0% and 42.8% to 63.5%, respectively; the core sets gave −6% and 63%. T8–T6 correlations used observed effects. This was one descriptive withdrawal series; the envelopes do not represent biological uncertainty, and post-withdrawal DNAm observations were not held out from feature selection.

A post hoc descriptive transfer applied the 123-CpG set and unchanged external centroid to HPF; all sites were present. Per-sample mean absolute reference distance was compared with the corresponding control. With one treatment/control pair per observation, no inferential test was applied. Deviation was higher than control at all eight observations (0.079–0.724 beta percentage points), including 0.253 at T6 and 0.692 at T8. This additional donor background does not provide replicated between-donor validation.

#### 8. Cancer preprocessing, reference scores and statistical families

TCGA duplicate sample identifiers were summed at raw-count level. For age analyses, pan-miR-29 summed miR-29a, both miR-29b loci and miR-29c before blind DESeq2 VST. Each cancer required at least 25 eligible tumours. The cohort comprised 8,084 primary and 344 metastatic samples from unique patients; breast contributed one metastasis and melanoma 343. RNA analysis excluded nonvariable features within cancer before repeated-subsample correlations. RNA and miRNA summaries weighted cancers equally. Age acceleration was the within-cancer residual of Horvath DNAmAge on chronological age, recomputed within each 80% subsample; miRNA was not separately age-adjusted in these correlations.

CCLE miRNA values came from the processed NanoString resource. Within each lineage, the three miR-29 measurements were standardised and averaged to form pan-miR-29. Chronological age denotes donor age. Acceleration was the residual of the specified clock prediction on age within the analysis cohort. The main rank used Skin & Blood acceleration; supplementary ranks additionally used donor age, Skin & Blood DNAmAge, Horvath pan-tissue DNAmAge and Horvath acceleration.

Paired analyses required primary tumour (TCGA type 01) and adjacent normal (11) from the same patient for both RNA and miRNA, with at least five complete pairs per histotype. Pan-miR-29 VST values were standardised over all tumour and normal samples within cancer, including unpaired samples. Each MRC gene was similarly standardised, the five-gene mean was calculated per sample, and the composite was standardised again. Effects were mean patient-level $Z_{tumour}-Z_{normal}$ differences, with two-sided one-sample t tests, 95% t intervals and BH correction across sixteen histotypes separately for each readout. Pooling used REML/Knapp–Hartung; the thirteen-organ summary first combined kidney and lung histotypes within organs. Paired-effect association used Pearson correlation across sixteen cancer means. Patient-level densities weighted each pair by the inverse of its cancer's pair count, giving equal total cancer weight. Between-histotype heterogeneity was approximately $I^2$ = 97% for both readouts.

Normal methylation references were defined after autosomal restriction, missingness filtering and KNN imputation. Own-reference eligibility required at least five strict methylation tumour–normal pairs and 5,000 stable CpGs. Reference medians were calculated on M values after beta clipping. Selection applied SD, age-correlation and age-slope bounds, without a sex-association filter. KICH lacked its own adjacent-normal methylomes and used 205 shared kidney normals; KIRC and KIRP used separate references. CHOL used nine normals. STAD had only two strict methylation pairs and was excluded from the fifteen-histotype reference-deviation cohort while remaining in paired-expression analyses.

The miRNA–reference-deviation analysis required primary-tumour methylation score, age, pan-miR-29 and a finite Horvath prediction; Horvath availability defined inclusion, not covariate adjustment. MRC analysis did not require miRNA or Horvath and therefore used a different overlap cohort. For reference-deviation associations, pan-miR-29 was calculated by summing DESeq2 size-factor-normalised abundances and applying $\log_2(x+1)$. Both variables were regressed on a natural spline of age with three degrees of freedom, followed by Spearman correlation of the residuals.

Cancer-level nominal tests used asymptotic Spearman tests (cor.test exact = FALSE); Fisher intervals used $SE_z = 1/\text{sqrt}(N-6)$. REML/Knapp–Hartung pooling produced fifteen-histotype and twelve-organ summaries, with inverse-variance weighting within organs at the first stage. Age-acceleration sensitivity analyses used BH correction across forty-five cancer-level tests (three models × fifteen histotypes) and separately across three pooled tests. Cancer and lineage omissions recomputed aggregate estimates as descriptive sensitivities.

Between-histotype heterogeneity was $I^2$ = 93.4% for miR-29 and 90.0% for MRC. Single-histotype omissions gave pooled correlations of −0.167 to −0.116 for miR-29 and 0.153 to 0.196 for MRC. A post hoc omission of BLCA and LUSC, the strongest miR-29-associated histotypes, gave $\rho = -0.087$ ($P = 0.057$) for miR-29 and $\rho = 0.138$ ($P = 0.004$) for MRC.

An additional model residualised miR-29 and reference deviation on the age spline and Horvath acceleration. The pooled association was $\rho = -0.150$ (95% interval −0.261 to −0.036; three-pooled-test BH q = 0.028), compared with $\rho = -0.145$ after age adjustment alone. Acceleration itself had a pooled association with reference deviation of $\rho = -0.0208$ (95% interval −0.1088 to 0.0676; q = 0.622; $I^2$ = 88.3%). This assessed covariate adjustment, not mediation.

Two post hoc cross-histotype comparisons distinguished mean reference deviation from the strength of its association with miR-29. The first related the mean paired decrease in pan-miR-29 to the mean paired increase in RMS M-value reference distance in the same patients. Pan-miR-29 used the existing VST-derived within-histotype standardised score, not a log fold change. Requiring at least five complete tumour–normal pairs with both assays retained 12 histotypes and 329 patients. COAD and LUSC had four and three complete pairs; KICH lacked own-normal methylation and STAD lacked the selected reference. Spearman rho was 0.140 (permutation P = 0.669; 95% bootstrap interval −0.580 to 0.770). Sensitivities using normal-SD-calibrated RMS distance, absolute beta distance, equal-organ aggregation and separate full paired cohorts gave rho = 0.280, 0.007, 0.191 and 0.174, respectively (all P ≥0.379). These comparisons did not establish that larger mean miR-29 decreases predict greater mean reference deviation.

The second comparison related each histotype's mean pan-miR-29 decrease from the original paired-expression cohort to the negative of its age-adjusted miR-29–reference-deviation correlation in Fig. 5h. A positive coefficient therefore indicates a stronger inverse association, not a greater amount of deviation. Across 15 histotypes representing 553 paired-expression patients, rho was 0.586 (P = 0.0233; 95% interval 0.090–0.874). Excluding KICH, which uses a shared kidney reference, gave rho = 0.490 (P = 0.0792; interval −0.043 to 0.825). All leave-one-histotype-out coefficients remained positive (0.490–0.705), although not all retained P <0.05. Both questions used equal-histotype weighting, two-sided tests with 100,000 permutations and plus-one correction, and percentile intervals from 20,000 histotype bootstrap resamples. BH correction across the two questions gave q = 0.669 and 0.0467. The second question was defined after the first result was known, before inspecting its own correlation. Histotype bootstrap intervals do not propagate all uncertainty in the source estimates. Neither analysis establishes patient-level prediction or quantifies an explanation of the heterogeneity in Fig. 5h.

**9. Public-resource provenance**

GTEx DNA-methylation profiles were obtained from GSE213478 [73].

The 113 developmental profiles were selected from GSE101571 (Wu et al.) [74], GSE157834 (the human component reported by Asami et al.) [75], GSE71318 (Dang et al.) [76], GSE65481 (Yanez et al.) [77] and GSE133854 (Leng et al.) [78]. The reprogramming mature-miRNA series GSE62776 was generated by Cacchiarelli et al. [79]. Cross-mammal methylation data came from the Mammalian Methylation Consortium, GSE223748 [80].

External reference selection used scalp/dura fibroblasts from GSE77135 [81], normal/control longitudinal cultures from GSE179847 [82], primary-cell subsets of GSE31848 [83] and purified leukocytes from GSE110554 [84]. Bulk-profile placement used the oral single-cell atlas GSE164241 [85].

**10. Software versions**

Local analysis environments used R 4.2.3. Expression and trajectory analyses used DESeq2 1.38.3, limma 3.54.2 and Slingshot 2.6.0. Statistical modelling used lme4 1.1-33, pbkrtest 0.5.2, emmeans 1.8.5, mgcv 1.8-42 and metafor 4.4-0 [86,87,88,89]; plotting used ggplot2 3.5.1. Network visualisation used

Cytoscape 3.10.3. Enrichment analyses used fgsea 1.24.0, and Skin & Blood predictions used meffonym 1.2.1, as detailed above. The recorded activity-inference workflow used DoRothEA 1.10.0, decoupleR 2.4.0, VIPER 1.32.0 and PROGENy 1.20.0. Primary-HGF RNA-age predictions used RNAAgeCalc 1.10.0; PASTA used version 0.0.0.9000 (commit 58bcc7a69ee97f2dc9e3623ac86538c251ddd498). Figure 2 analyses used rotl 3.1.0 and ape 5.7-1. The hTERT-HGF tAge prediction environment used tAge 1.1.0 preprocessing and scikit-learn 1.3.2 [90]. Read-processing and sequence-alignment software versions are specified in Methods and section 2.

Server-side methylation preprocessing used R 4.4.2 with SeSAMe and sesameData 1.24.0. Oral single-cell reference processing used Seurat 5.4.0, SeuratObject 5.3.0 and Harmony 1.2.4 in the same R environment. The CHIR99021 analysis additionally used DMRcate 3.2.1, limma 3.62.1 and minfi 1.52.1, with the EPIC v2 manifest and hg38 annotation packages 1.0.0.

Acquisition software documented in the retained instrument records comprised NextSeq Control Software 4.2.0.112, Real-Time Analysis 2.11.3 and iScan Control Software 4.0.0.147 and 4.2.1.729.
